# Septins promote breast cancer cell invasion in 3D collagen gels by influencing actin-based protrusion formation

**DOI:** 10.64898/2026.07.17.739166

**Authors:** Anouk van der Net, Klara Beslmüller, Nicole van Vliet, Margherita Tavasso, Myrthe Beerens, Ruben C. Boot, Pouyan E. Boukany, Erik H. J. Danen, Gijsje H. Koenderink

**Affiliations:** Delft University of Technology, Department of Bionanoscience, Kavli Institute of Nanoscience, Delft, 2629 HZ, The Netherlands; Leiden University, Leiden Academic Centre for Drug Research, Leiden, 2333 CC, The Netherlands; Delft University of Technology, Department of Chemical Engineering, 2629 HZ, The Netherlands

## Abstract

Septins are cytoskeletal proteins that contribute to essential cellular processes such as cell migration and cell division through interactions with the cell membrane and the cytoskeleton. High expression of septins is correlated with breast cancer malignancy and promotes cell invasion, but the molecular complexity of septin’s interactions has made it challenging to dissect the underlying molecular mechanisms. Here, we used a conditional knockout approach to deplete SEPT7 in the metastatic triple-negative breast cancer cell line Hs578T and examined the role of septin in 3D-matrix invasion of breast cancer cells. We show by spheroid assays that SEPT7 deletion strongly impairs breast cancer cell invasion into collagen gels. Additional single-cell migration studies using 3D collagen gels and microfluidic pillar devices that mimic the pores present in collagen matrices showed that SEPT7 expression regulates confined cell migration through control of cell shape and actin-based protrusions.

## Introduction

Septins form a family of guanosine triphosphate (GTP)-binding proteins that is increasingly recognized as an essential component of the cytoskeleton in mammalian cells^1^. Humans possess 13 septin genes referred to as SEPT1-12 and SEPT14^2^. Paralogous septins are categorized into four groups based on sequence homology, named the SEPT2, SEPT6, SEPT7 and SEPT3 groups^3^. Combinations of pairs of septins from each group assemble into palindromic hetero-oligomers in a defined linear sequence^4, 5^. In mammalian cells, both hexamers (with the general sequence SEPT2-6-7-7-6-2) and octamers (with the general sequence SEPT2-6-7-3-3-7-6-2) are found in cell-type-specific proportions^6, 7^. Each septin subunit can be exchanged with another from the same group (Kinoshita Rule^8^) and with different isoform variants. In particular SEPT9, which belongs to the SEPT3 group, has many isoforms which are differentially expressed in different cell types and in neoplasia^9^. The large variety of ensuing septin hetero-oligomer compositions may contribute to redundancy, but also to cell type-specific functions^10^. Each septin subunit has two interaction sites for binding adjacent septins: one formed by its GTP-binding domain (G-interface) and the other by its N- and C-terminal extensions (NC-interface). This enables the formation of hetero-oligomers that assemble end-to-end into nonpolar thin (4 nm) filaments via their SEPT2 subunits^11^. These filaments can in turn laterally pair through interactions of the C-terminal coiled coil domains of SEPT6 and SEPT7^12^. It is still poorly understood how cells regulate septin self-assembly and turnover dynamics, although post-translational modifications of septins^13^ and interactions with Borg proteins^14–16^ are known to be important.

Septin filaments in mammalian cells are often found in association with the plasma membrane, where they serve as scaffolds that reinforce the membrane, organize signaling receptors, and form diffusion barriers (reviewed in^17^). Septins can directly bind to lipid membranes through interactions with phosphoinositide lipids, in particular phosphatidylinositol 4,5-bisphosphate (PIP2)^18^. However, septins can also indirectly associate with the cell cortex through interactions with the actin cytoskeleton. Actin-associated septins have been found to regulate the structural stability and contractility of actin stress fibers in adherent cells^19–21^, the actin cortex in immune cells^22^, and the actin cortex in epithelial and endothelial tissues^23, 24^. Septins may also bind to microtubules through a short amino-terminal repeat motif unique to the septin 9 isoform SEPT9-i1^25, 26^. Recent work suggests that septin 9 phosphorylation can act as a molecular switch between actin and microtubules binding^27^.

Through their interactions with cell membranes and the cytoskeleton, septins contribute to a wide range of cellular functions including cell division^28^, immune cell migration^22^, and mechanotransduction^29, 30^. As a consequence, septin dysfunctions are associated with a wide range of diseases including different forms of cancer^31^. Some septin paralogs function as suppressors of tumor cell invasion. In glioma cells, for instance, high SEPT7 expression is linked to inhibition of cell invasion^32, 33^. In contrast, various septins function as tumor cell invasion promoters in breast cancer. For example, high expression of SEPT2, SEPT7 and SEPT9 in breast cancer is associated with more aggressive forms of the disease^34, 35^. High expression of SEPT9 is associated with poor clinical outcomes and resistance to cancer treatment in breast and other solid tumours^36, 37^. In various breast cancer cell lines, septin knockout or inhibition of septin filament dynamics with forchlorfenuron was shown to lower cell proliferation and invasion rates, while septin over-expression had the opposite effect^34–36^. SEPT9 overexpression was shown to promote metalloprotease-dependent invasion of breast cancer cells^38^.

It is challenging to dissect the mechanisms by which septins impact cancer cell invasion. Cancer cell invasion is a complex process that relies on all cytoskeletal networks jointly regulating cell deformability, cell contractility, integrin-mediated matrix adhesion, and the formation of specialized membrane protrusions. Although research till now has mainly focused on the roles of actin and microtubules, there is evidence that septins also participate in all of these functions. Septins can influence cell deformability by membrane and/or actin cortex reinforcement^39^. They influence cell contractility by regulating the organization and the activity of actin stress fibers through interactions with non-muscle myosin II and myosin binding partners^6, 20, 40–46^, influencing nuclear and genomic stability during cell squeezing^30, 47^. By stabilizing actin stress fibers, septins also promote integrin-mediated matrix adhesion^6, 48, 49^. In migrating endothelial cells, septins furthermore promote focal adhesion turnover by guiding growing microtubuli towards focal adhesions^50^. Finally, septins influence the formation of distinct types of membrane protrusions that drive cell migration. Septins stabilize actin-rich lamellipodia that drive mesenchymal migration, regulate blebs that drive amoeboid migration^22, 45, 51^, promote the maturation of podosomes that facilitate metalloprotease-dependent invasion^52^, and control the formation of microtubule-based microtentacles that promote metastatic colonization of circulating tumor cells^53^. Curvature-sensitive membrane binding may potentially contribute to the preferential association of septins with membrane protrusions^18, 54^ and subsequent reinforcement of membrane deformations^55^. In addition, there is evidence that septins influence protrusion formation through interactions with regulators of small G proteins that control actin polymerization and contractility^36, 56–60^.

Prior studies of septins in breast cancer cells mainly focused on 2D migration across flat substrates. Here, we instead investigate the role of septins in 3D invasion of collagen networks, with the aim to capture the confinement experienced by cancer cells during tissue invasion. We generated an inducible CRISPR/Cas9 knockout of SEPT7 in Hs578Ts cells, a metastatic triple negative human breast cancer cell line that is resistant to various therapeutic strategies^61^. Hs578T cells express SEPT1 and SEPT2 (both SEPT2 group), SEPT7, SEPT9, and SEPT11 (SEPT6 group), which co-localize with actin filaments and not with microtubules^53^. To knockout septins, we decided to target SEPT7, since this septin is the only member of its family and hence plays a critical role in the stability of septin hetero-oligomers. Whenever SEPT7 is silenced or knocked out, expression levels of other septins tend to go down as well and the remaining septins are unable to form functional hetero-oligomers or filaments^39^. We first investigated how SEPT7 depletion influences breast cancer cell invasion using 3D spheroid invasion assays in collagen hydrogels. Given the widespread role of septins in cytoskeletal reorganization and membrane dynamics described above, we hypothesized that septins influence 3D breast cancer cell invasion by affecting cell shape and specifically protrusion formation. To test this hypothesis, we quantitatively investigated the shape control of SEPT7 knockout cells in collagen gels and in rigid micropatterned environments (quasi 1D-micropatterns and 3D-constrictions). We found that SEPT7 knockout strongly impairs 3D cell invasion as well as membrane protrusion formation, indicating that SEPT7 is a crucial regulator of confined cell migration by promoting the formation of actin-rich protrusions required for crawling through the extracellular matrix.

## Results

### Characterization of SEPT7 depletion in Hs578T cells

To test the impact of septins on breast cancer cell invasion of 3D collagen matrices, we decided to create a conditional SEPT7 knockout model based on Hs578Ts cells, a metastatic triple negative human breast cancer cell line^61^. We generated doxycyclin-inducible Hs578T-Cas9 and transduced them with a short guide RNA (sgRNA) targeting SEPT7. Next, the cells were treated with doxycyclin to induce Cas9 expression and knock out SEPT7 expression (SEPT7 cKO). Control (NT CTRL) cells were obtained by performing the same treatment but using non-targeting sgRNA. We tested by immunofluorescence microscopy (Fig. S1) and Western Blot analysis (Fig. S2) at different induction times (2, 5, 6, and 7 days) that 7-day treatment with doxycyclin was optimal for efficient SEPT7 depletion.

To qualitatively characterize the efficiency of septin depletion, we first compared immunocytochemistry images of cells before doxycyclin treatment (day 0) and after 7 days of doxycyclin treatment, staining for SEPT7 and F-actin. As expected, in control cells, SEPT7 was clearly visible as linear signals co-localizing with actin stress fibers (Fig. 1A(i-iii)). After 7 days of doxycyclin treatment, an estimated 80-90% of the cell population lost visible SEPT7 expression (Fig. 1A(iv-vi)). Quantitative analysis by Western Blot analysis showed that doxycyclin treatment reduced the expression level of SEPT7 on average by 6-fold (Fig. 1B(i)). As SEPT7 is critical for the stability of septin hetero-oligomers^4, 6^, we also expected a decrease in the levels of other septins. Western Blot analysis indeed also showed markedly reduced expression levels of both SEPT2 (∼ 1.3-fold decrease) and SEPT9 (∼ 2.4-fold decrease) after doxycyclin treatment (Fig. 1B(ii-iii)). Immunocytochemistry images confirmed that SEPT7 cKO cells also had a markedly lower signal for SEPT9 as compared to control cells (Fig. 1C(i-ii,iv-v)). We also found that the SEPT7 cKO cells regularly exhibited a disrupted Golgi apparatus (Fig. 1C(iii,vi)) and dysregulated microtubules with dense compact structures near the cell cortex (Fig. 1D(ii,iv)). Both of these aberrant structures co-localized with residual SEPT9 structures (Fig. 1D(iii)). Western blot analysis of Hs578T cells did not show any change in the level of α-tubulin upon SEPT7 depletion (Fig. S10E).

**Figure 1.**
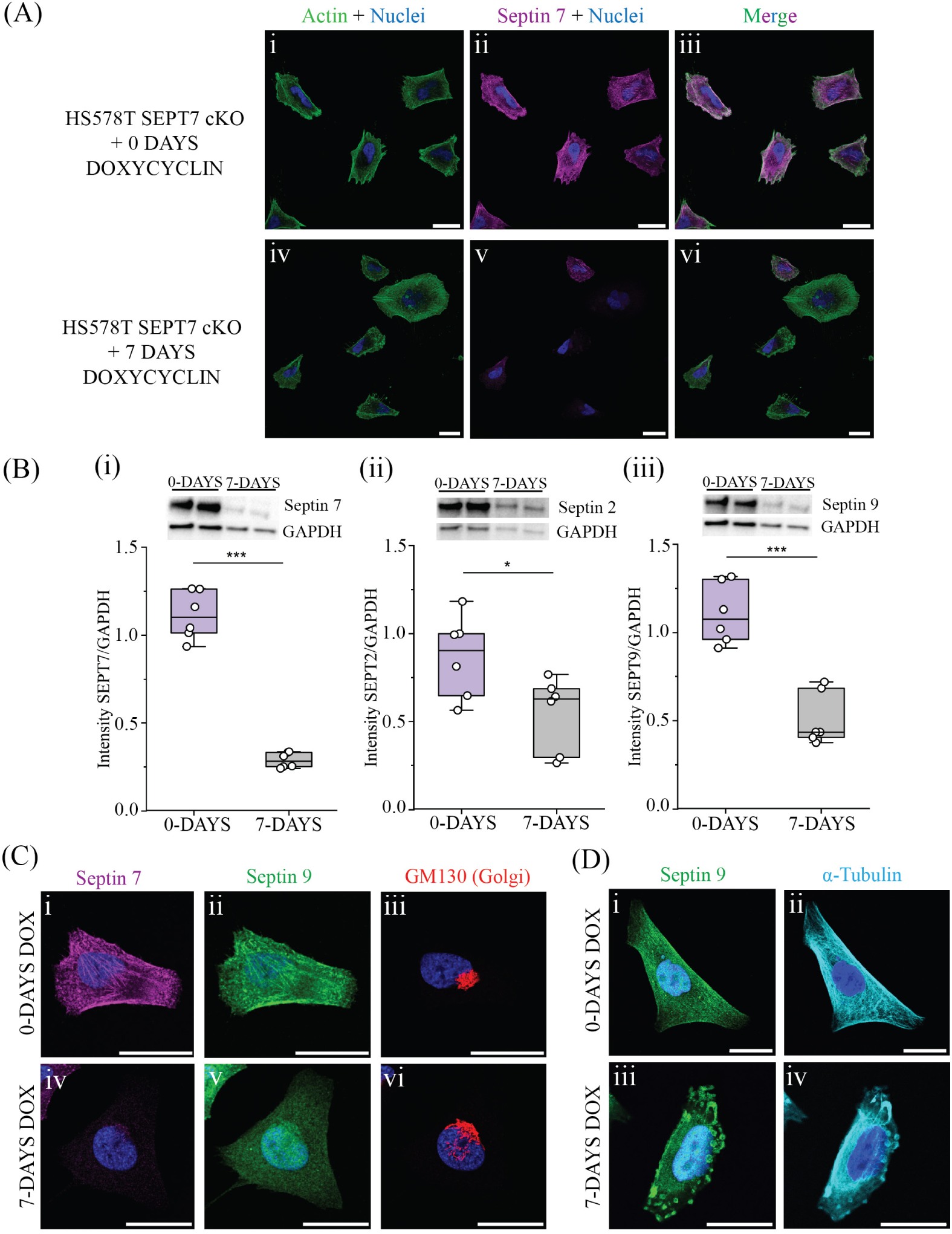
Characterization of inducible septin 7 (SEPT7) knockout in Hs578T breast cancer cells. (A) Immunocytochemistry images and Western Blot quantification of SEPT7 expression in Hs578T SEPT7 cKO cells at day 0 (panels i-iii) and day 7 (panels iv-vi) of doxycyclin treatment. Cells are stained for F-actin (green), SEPT7 (magenta) and nuclei (blue). Scale bars are 50 µm. (B) Western Blot (top) and corresponding boxplots quantifying Western Blot intensities relative to GAPDH (bottom) for SEPT7 (panel i), SEPT2 (panel ii) and SEPT9 (panel iii) before and after 7 days of doxycyclin treatment. Data for intermediate time points are shown in Fig. S1. (C, D) Immunocytochemistry images of cells before (top rows) and after (bottom rows) 7 days doxycyclin treatment for SEPT7 (magenta), SEPT9 (green), GM130/Golgi (red), α-tubulin (cyan) and nuclei (blue). Scale bars are 25 µm. In the boxplots, (*) = p < 0.05, (***) = p < 0.001.

### Septins modulate actin organization and cell shape on 2D- and quasi 1D-substrates

In control cells, we observed that septins localize with actin stress fibers (Fig. 1A(iii)) but not with microtubules (Fig. 1D(i,ii)), consistent with previous findings for Hs578T cells^53^. Western blot analysis showed that SEPT7 depletion was associated with a slight upregulation of actin expression (Fig. S10A) while anillin levels did not change (Fig S10B). To investigate the interplay between actin and septin organization in Hs578T cells, we analyzed how each responded to depletion of the other. When we depleted actin fibers by cytochalasin D (CD) treatment, septins transformed from linear structures co-aligned with long actin stress fibers (Fig. 2A, top row; zoom-in areas indicated by white arrows) into shorter fibers, often crescent- or ring-shaped (Fig. 2A, middle row). Conversely, SEPT7 depletion from the cells resulted in minor changes in the organization of the actin cytoskeleton (Fig. 2A, bottom row). The most prominent change was a clear loss of actin fibers in perinuclear areas of the cells (areas indicated by white squares in Fig. 2C, left, zoomed-in images in Fig. 2C, middle and right): perinuclear stress fibers are visible in control cells (top row) but are lost in SEPT7 knockout cells (bottom row). In addition, control cells showed frequent linear actin-filled membrane protrusions, whereas SEPT7-depleted cells showed much fewer such protrusions (Fig. 2D, zoom-in areas indicated by white squares). Neither actin depolymerization nor SEPT7 depletion significantly impacted the spread areas of the cells (Fig. 2B). However, simultaneous actin depolymerization and SEPT7 depletion by performing CD treatment of SEPT7-cKO cells caused a significant reduction in cell area accompanied by anomalous cell shapes (Fig. S3). Thus, the actin and septin cytoskeleton jointly control cell spreading on flat 2D substrates.

**Figure 2.**
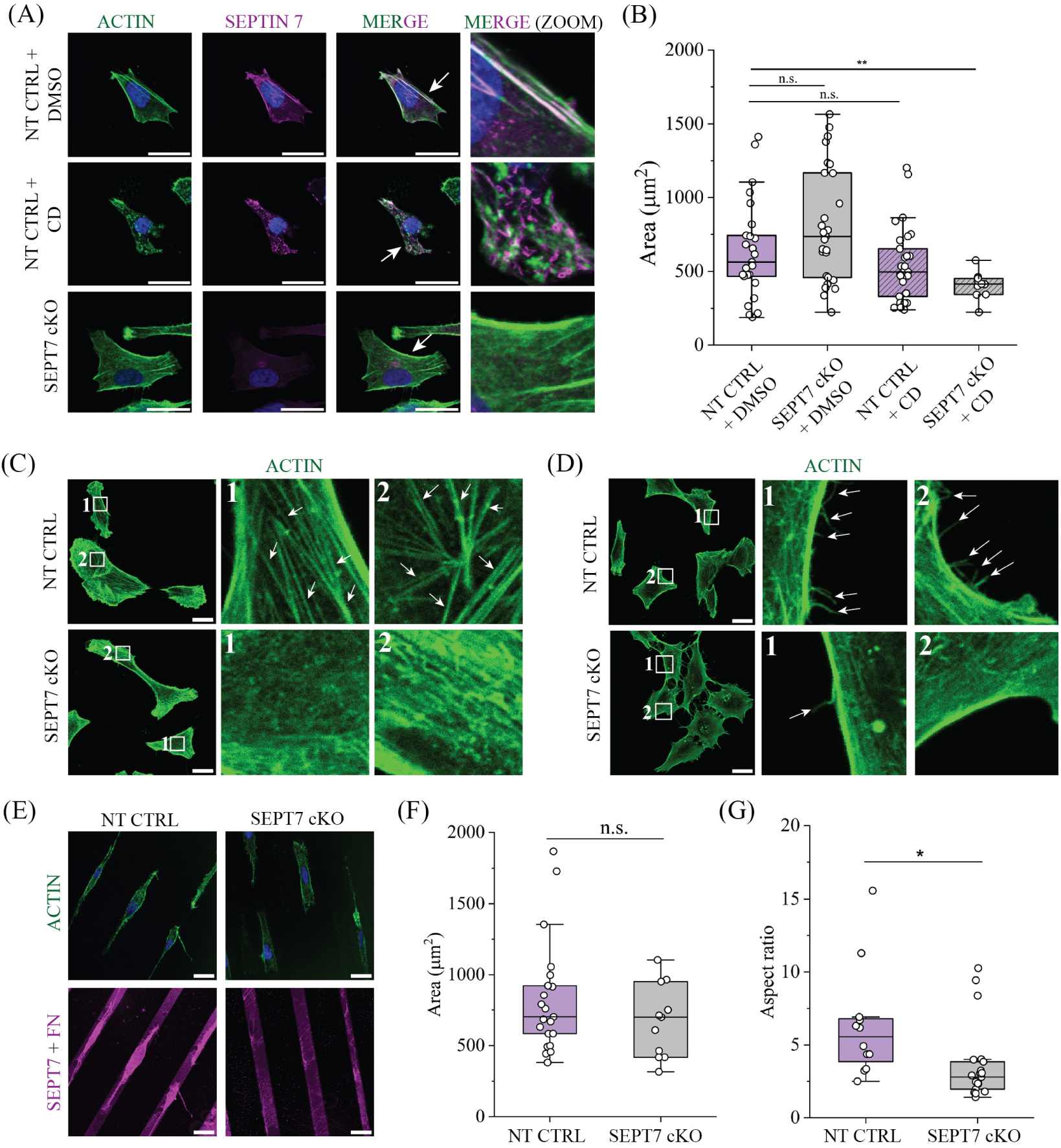
Interplay between septin and actin in Hs578T breast cancer cells. (A) Immunocytochemistry of control cells (NT CTRL) treated with vehicle control (DMSO, first row) or cytochalasin D (CD, second row) and SEPT7 knockout cells (SEPT7 cKO, third row). The columns show fluorescence signals of filamentous (F-)actin (green), SEPT7 (magenta) and nuclei (blue). White arrows in MERGE images indicate areas used for MERGE (ZOOM) images. (B) Boxplot showing the cell areas of control cells (NT CTRL + DMSO), cells depleted of SEPT7 (SEPT7 cKO), cells treated with CD to depolymerize actin (NT CTRL + CD), and SEPT7 cKO cells treated with CD (SEPT7 cKO + CD). (C) Immunocytochemistry images of F-actin in NT CTRL cells (top) and SEPT7 cKO cells (bottom). Numbered squares indicate zoomed-in areas for analysis of perinuclear regions. Stress fibers are indicated by white arrows. (D) Immunocytochemistry images of F-actin in NT CTRL cells (top) and SEPT7 cKO cells (bottom). Numbered squares indicate zoomed-in areas for analysis of the cell cortex. Membrane protrusions are indicated by white arrows. (E) Immunocytochemistry images of F-actin (top) and SEPT7 (bottom) for NT CTRL and SEPT7 cKO cells on micropatterned fibronectin-coated lanes with a width of 15 µm. Note that fibronectin is rhodamine-labeled and is therefore visible outside the cells in the SEPT7 channel. (F) Boxplot showing cell areas of micropatterned NT CTRL and SEPT7 cKO cells. (G) Corresponding boxplot of cell aspect ratios. In boxplots, (*) = p < 0.05, (**) = p < 0.005, (ns) = non-significant. All scale bars are 25 µm.

In the tissue microenvironment, breast cancer cells adhere to collagen fibers, which constrain cell adhesion to quasi-1D^62^. To test how septin depletion affects cell spreading under quasi 1D-confinement conditions, we seeded cells on top of 15 µm-wide micropatterned lanes coated with fibronectin. Control cells adapted to the pattern by forming rather regular elongated shapes with tapered ends and actin stress fibers running along their length (Fig. 2E, left). SEPT7 cKO cells showed notably less well-defined shapes and a less well-organized actin cytoskeleton (Fig. 2E, right). Quantification of the cell shapes showed that the SEPT7 cKO cells indeed had significantly smaller aspect ratios than control cells (Fig. 2G) despite having similar cell areas (Fig. 2F). We conclude that in Hs578T cells, SEPT7 interacts with the actin cytoskeleton and regulates perinuclear and cortical actin networks that are important for cell shape control under quasi-1D and 2D adherent conditions.

### Spheroid invasion in 3D collagen hydrogels is reduced in SEPT7 cKO Hs578T spheroids

Having established the Hs578T septin cKO model system, we turned to 3D invasion assays. We grew Hs578T cell spheroids and found that septin depletion reduced spheroid packing, elasticity and viscosity (Fig. S5 and Fig. S7). To create a physiologically relevant context, we embedded Hs578T cell spheroids in 2.4 mg/mL collagen gels, providing confined conditions (average pore size ∼1.3 µm^63^). We performed live-cell imaging of the spheroids to track the progression of cell invasion. At the start (defined as t = 0 h, circa 2 hours after embedding the spheroid in collagen), we already observed distinct differences in the number of protrusions exhibited by control versus SEPT7 knockout spheroids (Fig. 3A). NT CTRL spheroids showed clear protrusive behavior into the collagen gel at various sites along their contour, while SEPT7 cKO spheroids had a limited number of protrusions within the same incubation time.

**Figure 3.**
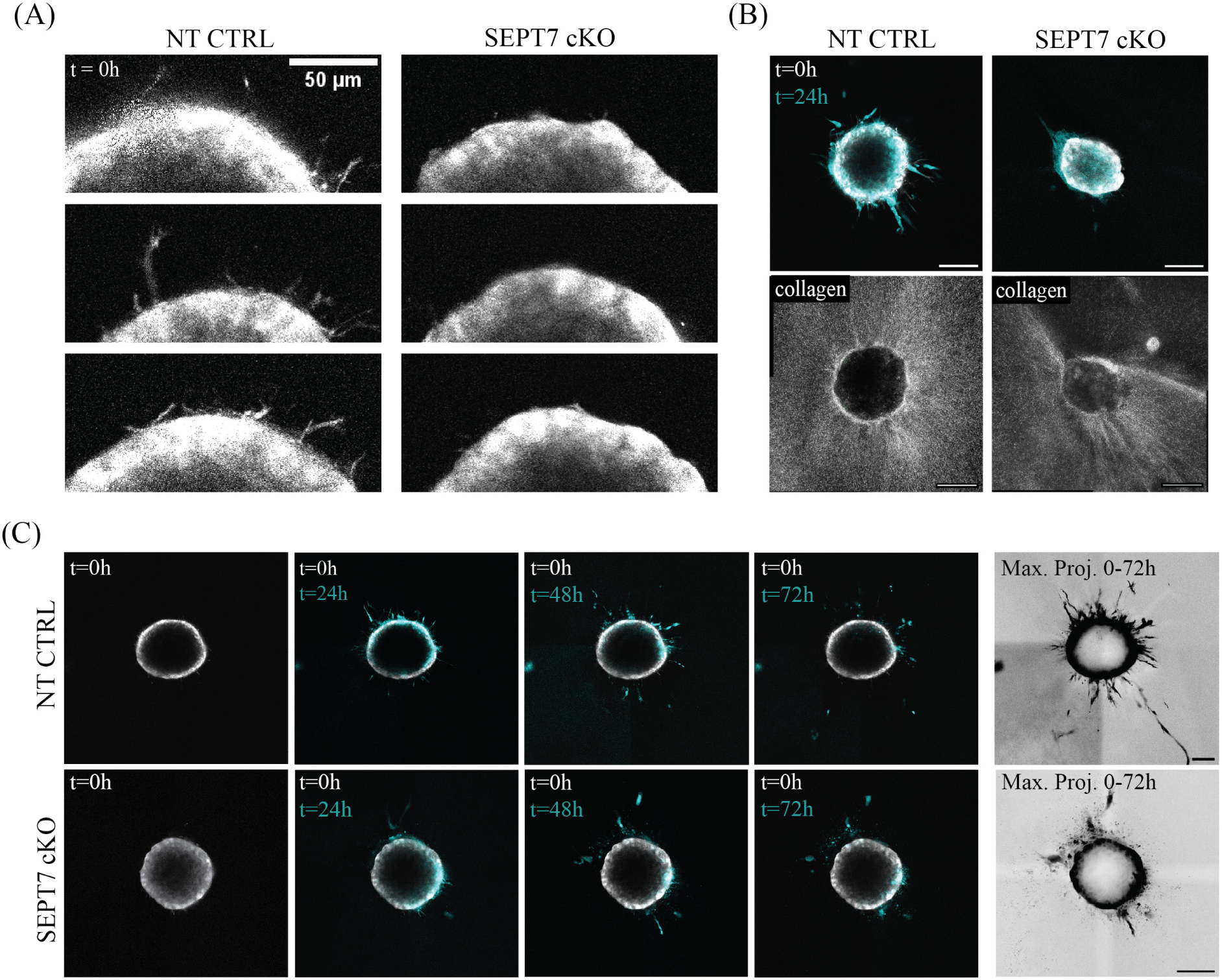
Impact of SEPT7 knockout on Hs578T cell invasion in fibrillar collagen type I networks measured by live-cell confocal imaging of 3D spheroid invasion. (A) Spheroid protrusions stained with Cytotracker Orange (gray) imaged at t = 0 h (∼2 hours after placing spheroid in 2.4 mg/mL collagen) for NT CTRL (left) and SEPT7 cKO (right) cells. Images are taken at the spheroid equators (rows show different replicates, n=3). (B) Correlation between cell invasion and collagen network remodeling. Top panels show an overlay of spheroid invasion measured by the Cytotracker Orange signal at t = 0h (gray) and t = 24h (cyan), showing the extent of invasion for NT CTRL (left) and SEPT7 cKO (right) spheroids at the spheroid equators. Bottom panels show the corresponding reflection images of the collagen network at t = 24h of invasion. (C) Time lapse image series comparing invasion of NT CTRL (top) and SEPT7 cKO (bottom) spheroids at the equators. First four columns show overlays of Cytotracker Orange signal at different time points (t = 24h, t = 48h and t = 72h, cyan) with the initial image at t = 0h (gray). Far-right columns show maximum time projections (constructed from maximum intensity projections) of the spheroid invasion from 0 - 72h. Scale bars are 100 µm. N = 6-9 for 24h live-cell experiments and N = 3 for 72h live-cell experiments.

Imaging the invasion process over a period of 24 hours revealed that also at later times, 3D invasion was noticeably impaired by septin depletion. NT CTRL spheroids consistently protruded in more areas and further away from the original spheroid border than SEPT7 cKO spheroids (Fig. 3B). In case of NT CTRL spheroids, we observed mesenchymal-like invasion of single cells that detached from the spheroids. Interestingly, SEPT7 cKO cells often invaded in a distinct way, where only cells on two opposite ends of the spheroid invaded in a collective manner, with little to no cell detachment (see Fig. 3B, top right panel). The resulting ’polarized’ spheroid invasion shape also affected the collagen network surrounding the spheroids, with anisotropic collagen fiber alignment around SEPT7 cKO spheroids compared to more isotropic collagen fiber alignment surrounding NT CTRL spheroids (see Fig. 3B, bottom panels).

To test whether these differences persist on longer time scales, we also performed live-cell imaging of spheroid invasion for 72 hours (Fig. 3C). Consistent with the 24 hour invasion assays, SEPT7 depletion again strongly restricted cell invasion, indicated by invasion in fewer areas and less far from the spheroid. In the example in Fig. 3C, NT CTRL cells reached distances of more than 500 µm from the spheroid edge, while SEPT7 cKO cells reached a maximal distance of ∼120 µm.

### SEPT7 is necessary for 3D spheroid invasion

To evaluate whether the invading SEPT7 cKO cells still contain residual SEPT7, we analyzed the SEPT7 expression of cells within the spheroids by immunocytochemistry. As expected, invaded cells in the NT CTRL spheroid assays showed consistent SEPT7 expression (Fig. 4, left). Interestingly, we also observed some residual SEPT7 expression in the SEPT7 cKO assay, but this was restricted to the few cells that protruded into the collagen network (Fig. 4, right). This observation suggests that cells with residual SEPT7 expression are the ones that are responsible for the residual invasion in the SEPT7 cKO assays. Together with the live-cell imaging data showing that invasion is strongly impaired for SEPT7 cKO spheroids, this indicates that SEPT7 expression is essential for 3D spheroid invasion of Hs578T breast cancer cells.

**Figure 4.**
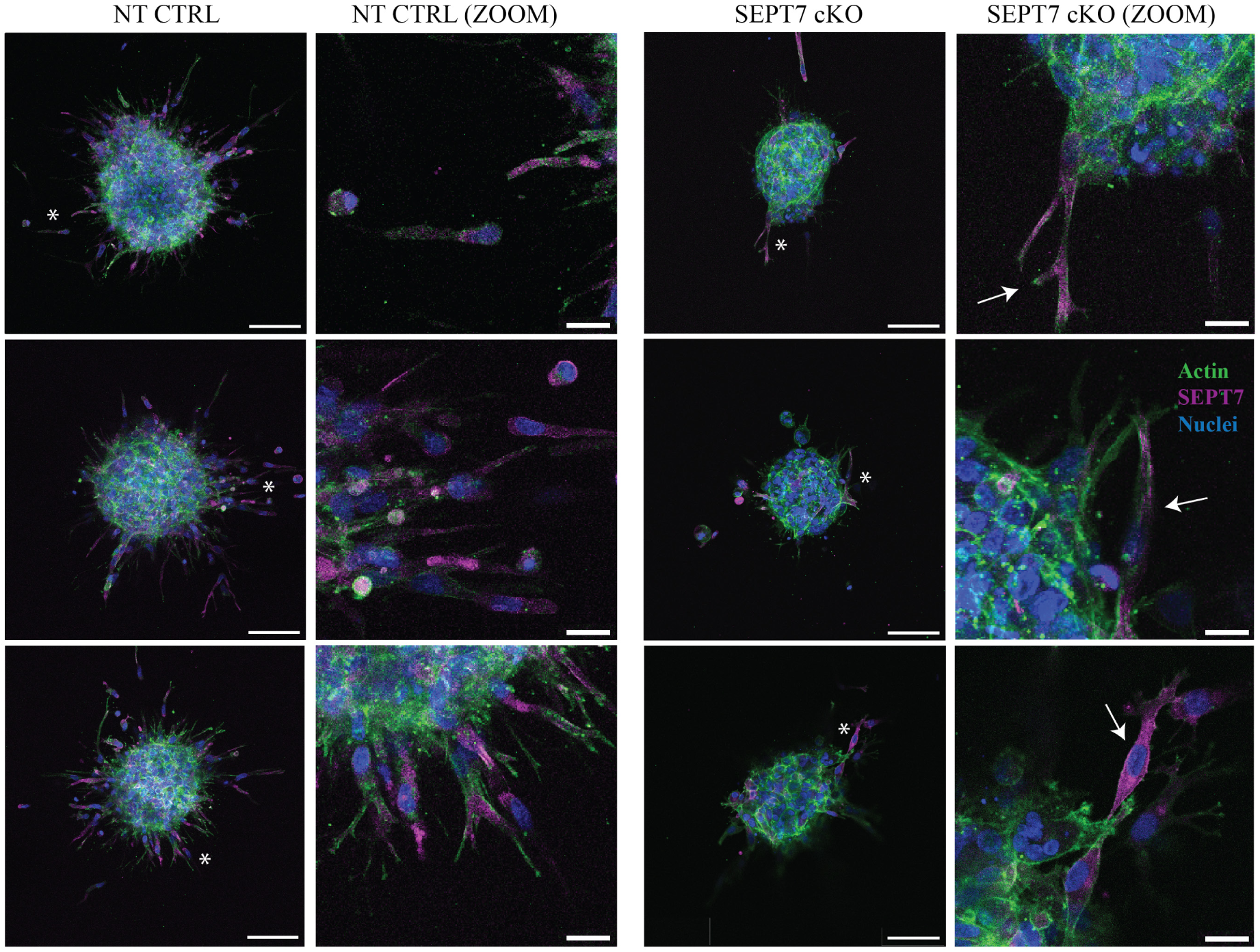
Immunofluorescence imaging of spheroids in collagen networks shows that invading Hs578T cells contain SEPT7. NT CTRL spheroids (left) and SEPT7 cKO spheroids (right) imaged after 24h of invasion in 2.4 mg/mL collagen gels, stained for actin (green), SEPT7 (magenta) en nuclei (blue). Rows show different replicates. Images are maximum projections of 70 µm Z-stacks with 10 µm Z-steps, with the spheroid equator positioned in the middle of the Z-stack. White asterisks on spheroid images indicate areas used for zoom-in (ZOOM) images. White arrows in SEPT7 cKO zoom-in images indicate invaded cells with residual SEPT7 expression. Scale bars are 100 µm for whole-spheroid images and 25 µm for zoom-in images. N = 6-9 for each condition, consisting of three biological replicates with 2-3 spheroids each.

### Septins govern 3D invasion potential and directional persistence of Hs578T cells

Time-lapse imaging of spheroid invasion showed that Hs578T cells confined in 2.4 mg/mL collagen networks exhibit a mesenchymal migration phenotype where cells invade individually. We therefore decided to quantify the invasion potential of individual control and SEPT7 knockout cells into collagen. To this end, we plated the cells on non-coated glass at sub-confluent density and tested what percentage of cells were able to migrate upwards into the 3D collagen network layered on top of them within a period of 3 days (Fig. 5A). To count the cells, we imaged the cell nuclei in the first 250 µm of the collagen network using Hoechst labeling.

**Figure 5.**
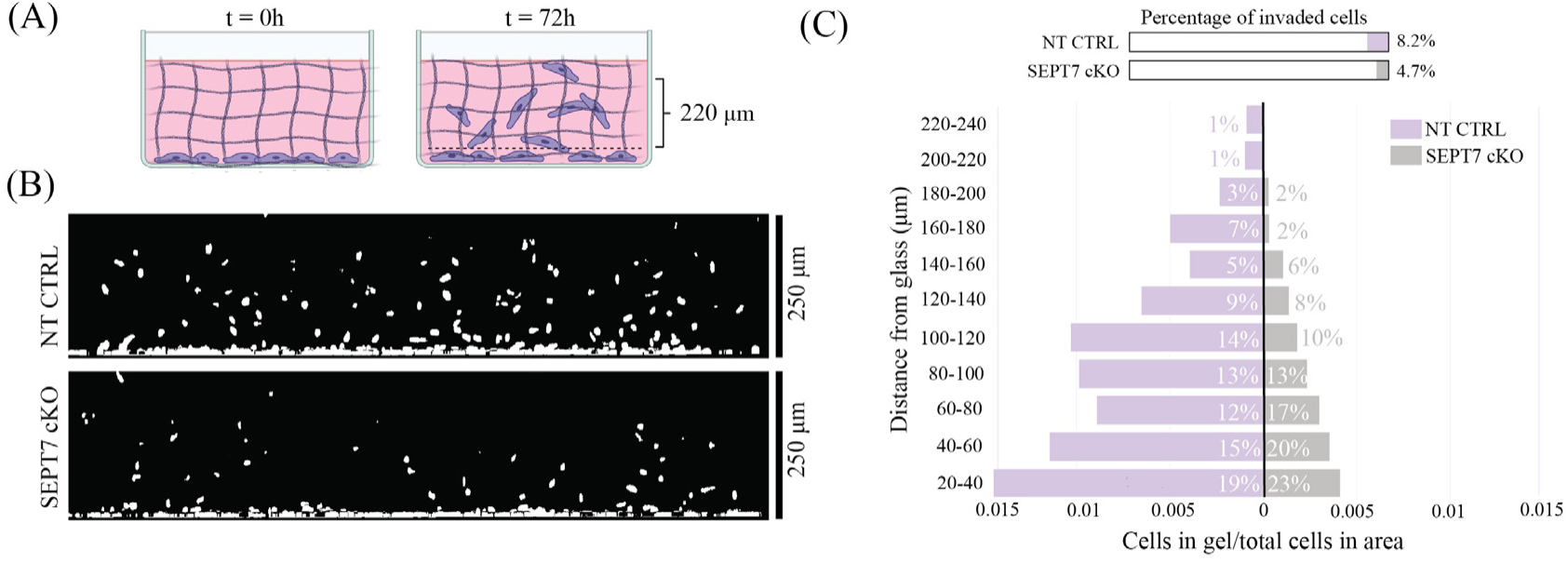
Quantitative analysis of the effect of SEPT7 knockout on single-cell invasion into collagen networks. (A) Schematic of the assay. The cells are seeded on glass and overlaid with a 2.4 mg/mL collagen network at t=0 hr (left). At t=72 hr, we image a 250 µm confocal z-stack and count the cells remaining on the glass and those that invaded into the collagen gel (right). (B) Side views (XZ-projections) of 250 µm Z-stacks showing the nuclei (Hoechst label) of NT CTRL cells (top) and SEPT7 cKO cells (bottom). Cells localized within 0-20 µm of the glass are classified as non-invaded. Cells localized between 20-240 µm (220 µm total) are counted as invaded. (C) Quantification of the percentage of invaded cells shown as a bar diagram (top) and as a population pyramid (bottom) demonstrates differences in the extent and range of invasion for NT CTRL (purple) and SEPT7 cKO (gray) cells. Percentages inside bars show the distribution of invaded cells, binned for 20 µm Z-steps. For ctrl conditions, two biological replicates were imaged at two different areas, resulting in n = 4. For knockout conditions, three biological replicates were imaged at two different areas, resulting in n = 6.

Side views (XZ-projections) of the invaded collagen gels indicate impaired invasion of SEPT7-deficient cells (Fig. 5B). To quantify the invasion potential, we measured the number of cell nuclei observed within the Z-range of 20 - 250 µm and normalized this by the total cell count in these areas (over the entire Z-range of 0-250 µm). As shown in Fig. 5C, we indeed measured a larger percentage of invaded cells for NT CTRL cells (8.2%) as compared to SEPT7 cKO cells (4.7%). Moreover, binning the distribution of invaded cells in 20 µm Z-steps showed that the SEPT7 cKO cells invaded the collagen networks significantly less far compared to NT CTRL cells, with cells detected up to 200 µm versus 250 µm, respectively. These results demonstrate that SEPT7 depletion reduces both the invasion potential and the directional persistence of breast cancer cells.

### Septins promote cell elongation and protrusion formation during confined migration

Since 3D mesenchymal migration requires a distinct polarized cell shape with lamellipodial-like protrusions in front, we tested whether SEPT7 depletion affects the shape of individually migrating Hs578T cells in collagen networks. To this end, we seeded NT CTRL and SEPT7 cKO cells in collagen networks of the same concentration (2.4 mg/mL) as used in spheroid assays. We immunostained the cells after 48 hours for nuclei, actin and SEPT7. Confocal imaging revealed a striking difference in cell shape, with NT CTRL cells showing much more complex shapes than SEPT7 cKO cells (Fig. 6A). Control Hs578T cells showed a variety of shapes with numerous protrusions at the leading edge (left column in Fig. 6A, marked NT CTRL). In addition, they often appeared in a ’crab’ shape with the nucleus positioned in the middle, from which two equally sized ’arms’ protruded forwards (see middle and bottom panels in Fig. 6A, NT CTRL). Confocal reflection imaging revealed that the cells interacted with the collagen matrix, as indicated by aligned collagen near protrusions. Septin depletion led to a marked change in cell shape, with fewer protrusions and more compact configurations (right column in Fig. 6A, marked SEPT7 cKO). Confocal reflection imaging showed that the SEPT7 cKO cells still interacted with collagen, but only at the cell poles, where collagen alignment is visible.

**Figure 6.**
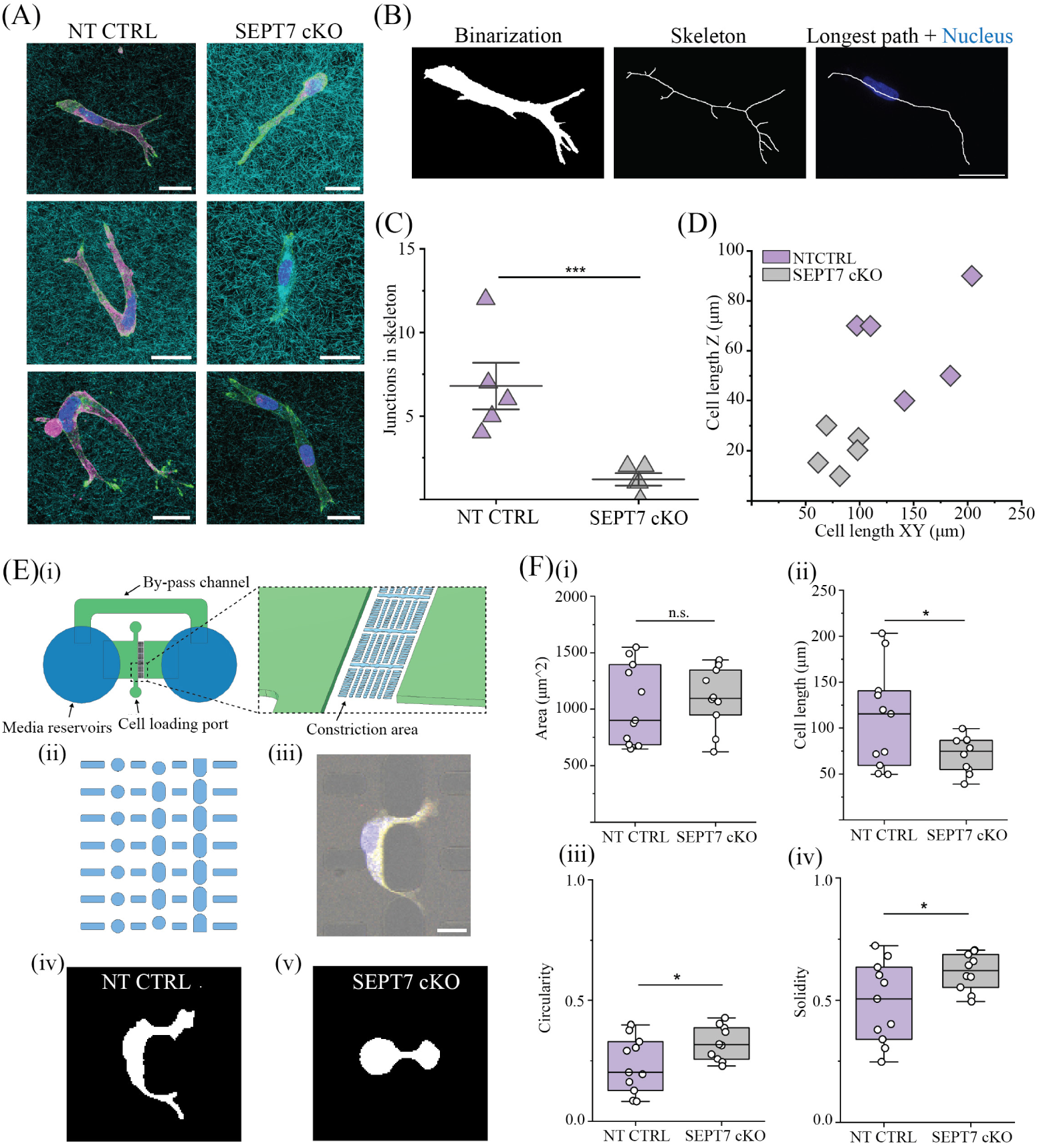
Impact of SEPT7 knockout on the shape of cells performing confined migration in collagen matrices or in matrix-mimicking microfluidic constrictions. (A) Maximum intensity Z-projections (Z-steps of 0.25 µm, Z-stack size dependent on cell size, ranging from 8.5 µm - 52.5 µm) of confocal immunofluorescence images of NT CTRL cells (left) and SEPT7 cKO cells (right) in a 2.4 mg/mL collagen gel after 48 hours. Cells were stained for nuclei (blue), actin (green) and SEPT7 (magenta), while collagen (blue) was imaged by reflection microscopy. Scale bars are 25 µm. Corresponding separate panels are shown in Fig. S4. (B) Image analysis pipeline for cell shape analysis. Maximum Z-projections were binarized (left) and used for skeleton detection (middle), from which we determined the cell length (longest path) and number of junctions (right). (C) Scatter interval plot of the number of junctions counted in each skeleton from Z-projections of NT CTRL (magenta) and SEPT7 cKO (gray) cells. (D) Scatter plot for the detected cell lengths in XY-plane and along Z-axis in 3D stacks of NT CTRL (magenta) and SEPT7 cKO (gray) cells. (E) (i) Schematic of the microfluidic set-up with zoomed-in schematic of the constriction area. (ii) The constriction area is designed as an array of pillars. (iii) Microscopy image of a Hs578T NT CTRL cell migrating in between the pillars, constructed from bright-field (gray) and fluorescent channels for cytoplasm (yellow) and nuclei (blue). Scale bar is 25 µm. (iv-v) Binarized images from cells in the constriction area, representative of NT CTRL (iv) and SEPT7 cKO (v) shapes during constriction. (F) Cell shape parameter boxplots, measured from NT CTRL (magenta) and SEPT7 cKO (gray) binarized images, for (i) cell area, (ii) cell length, (iii) circularity and (iv) solidity. (***) = P<0.001, (*) = P<0.05, (n.s.) = non-significant.

To quantify the difference in protrusion formation by control versus SEPT7 cKO cells, we performed automated skeleton detection and counted the number of junctions in the skeletonized cells as an indicator for cell shape complexity (Fig. 6B). As shown in Fig. 6C, SEPT7 depletion significantly decreased the complexity of the 3D cell shapes as compared to control cells, indicative of less (prominent) protrusions. The SEPT7 cKO cells were also consistently less elongated in both the XY-plane and along the Z-axis compared to control cells, as shown by analyzing the cell lengths in all three dimensions (Fig. 6D). These results indicate that SEPT7 contributes to cell shape regulation during confined 3D cell migration, promoting protrusion formation and cell elongation.

We hypothesized that the breast cancer cells need protrusions to navigate the geometrically complex pore space presented by the collagen gels. To test this idea and examine the impact of SEPT7 depletion on navigation of narrow pores, we seeded NT CTRL and SEPT7 cKO cells in a microfluidic device with rigid pillars providing constrictions that mimic the interconnected narrow pores present in collagen matrices (Fig. 6E(i-ii)). We imaged the cytoplasm and nucleus of cells within the constrictions to analyze the cell shapes during confined migration (Fig. 6E(iii)). Similar to cells in collagen gels, we found that SEPT7 cKO cells migrated with rounder and less complicated shapes compared to NT CTRL (Fig. 6E(iv-v)). In contrast, NT CTRL cells more often showed protrusions that entered one or multiple constrictions, indicating a ’probing’ shape (Fig. 6E(iv)). These findings have a striking analogy to the 3D cell shapes presented in the collagen gels. Depletion of SEPT7 did not affect average cell area (Fig. 6F(i)), but decreased cell elongation (Fig. 6F(ii)), and increased cell circularity and solidity (Fig. 6F(iii,iv)) during squeezing. These quantifications further demonstrate the loss of complexity from the cell shapes upon SEPT7 depletion. Altogether, we find that SEPT7 is necessary for the formation of cell protrusions and probing phenotypes during confined migration, both in microenvironments with regular and more irregular pore spaces.

## Discussion

Septins are cytoskeletal proteins that assemble into higher-order structures and interact with various key cellular components to support functions like cell polarity and cell shape control. High septin expression was previously correlated with metastatic breast cancers^34–36, 38^, but the exact mechanisms by which septins contribute to the enhanced invasion potential of breast cancer cells remain unknown. By using a gene-edited breast cancer model based on the metastatic triple negative human breast cancer cell line Hs578T together with a series of 2D and 3D microenvironments that mimic tissue confinement, we showed that SEPT7 expression regulates the invasion potential of Hs578T cells by allowing the formation of actin-based protrusions needed for navigating the collagen interstitial matrix.

Metastasis is driven by reorganization and crosstalk between different cytoskeletal networks. Because septins are known to influence the actin and microtubule cytoskeleton and their crosstalk, we first characterized the effects of SEPT7 depletion on the architecture of the cytoskeleton in 2D-adherent cells. Depletion of SEPT7 (which was accompanied by SEPT2 and SEPT9 depletion) resulted in a loss of peri-nuclear actin stress fibers and filopodial actin protrusions. The loss of peri-nuclear stress fibers could reflect destabilization due to the loss of septin-mediated scaffolding. Alternatively, there could be a disturbed balance between different SEPT9 isoforms in response to SEPT7 depletion, as Sept9i2 expression is associated with a loss of perinuclear stress fibers^64^. The organization of the septin and actin cytoskeleton were interdependent, since actin depolymerization by cytochalasin D caused septins to form rings, consistent with observations in other cell lines^19, 65^. We found that SEPT7 depletion also resulted in dysregulated Golgi and microtubule networks co-localized with residual SEPT9. We suspect that these defects arise from abnormal SEPT9 localization, since SEPT9 has previously been shown to be essential for regulating the integrity of the Golgi apparatus^66, 67^.

In SEPT7 cKO cell cultures we also noticed an increase in giant and multi-nucleated cells (Fig. S9A,B), in line with previous reports of septin-depleted human breast cancer cells^34^. However, the percentage of these giant cells in the cultures was only 5 - 10 % and, qualitatively, proliferation at a population level was not greatly affected. In many cell types, septins have been shown to be necessary for cell division through their interactions with the contractile actomyosin ring and abscission machinery^68^. However, some cell types such as hematopoietic cells can switch to septin-independent mechanisms of division^69^. Our observations suggest that the Hs578T cells also have ways to overcome septin-deficiency. Interestingly, we observed examples of cells that appeared to divide by crawling away from each other (Fig. S9C,D). This mechanism of cytokinesis closely resembles cytofission observed in giant multinucleated amoebae, which is driven by cortical actin waves^70^. This mechanism could be mediated by increased contractility in giant cells, as immunocytochemistry of phospho-myosin light chain 2 showed an increased signal in giant SEPT7 cKO cells (Fig. S11B). This increase was not accompanied by significant changes in phospho–myosin light chain 2 or myosin light chain protein levels as measured by Western Blot analysis (Fig. S11A), indicating that the increase does not occur across the entire SEPT7 cKO population.

To test the impact of SEPT7 knockout on Hs578T cell invasiveness, we performed 3D spheroid invasion assays in collagen networks. Upon SEPT7 knockout, we observed a strong inhibition of cell detachment and invasion from spheroids over a 72 hours time frame. Immunofluorescence microscopy showed that residual invasion originated from a small fraction of remaining SEPT7 positive cells within the knockout culture. Moreover, septin cKO spheroids tended to show collective invasion, in marked contrast to single-cell invasion observed for control spheroids. It will be interesting to further investigate the molecular basis of this change in migration mode, since collective invasion is related to increased metastatic site formation^71^. We propose that the heterogeneity of the knockout spheroids, which consist of mainly immobile (SEPT7-depleted) cells and a small fraction of motile cells (with residual SEPT7), may impact the spheroid unjamming transition^72^. We noticed that the edges of SEPT7 cKO spheroids before embedding in collagen appeared more rough than for control spheroids, which could indicate a lower packing density and reduced cohesion of the spheroid (Fig. S5). To test this idea, we measured the viscoelastic properties of the spheroids in a microfluidic device designed for spheroid compression^73^ (Fig. S7A,B). We found that SEPT7 depletion significantly decreased spheroid elasticity (Fig. S7C), which could be a direct result of septin loss or an indirect result from loss of actin stress fibers. Furthermore, our measurements indicated a trend towards lower viscosity (Fig. S7D), which could be related to lower cell packing densities. Analysis of the effect of SEPT7 knockout on cell-cell interactions and the deposition of extracellular matrix proteins within spheroids, for instance by Western Blot analysis, could further elucidate the role of septins in spheroid viscoelasticity, a factor that may impact the mode of invasion when the spheroids are embedded in collagen.

Since we observed mainly single-cell mesenchymal migration for Hs578T spheroids in collagen, we next performed single-cell 3D collagen migration assays to test how SEPT7 depletion impacts cell shape regulation in 3D-confined conditions. We found that SEPT7 depletion impaired cell invasion in 3D collagen gels, in line with the observations from the spheroid assay. Furthermore, SEPT7 depletion strongly impacted cell shape regulation. The control cells adopted crab-like shapes, extending long arm-like protrusions along different migration paths. Recent theoretical work showed that cells adopt these branched shapes in complex environments where their path branches into several alternative directions of migration^74^. Actin polymerization at the leading edge generates multiple competing protrusions to choose the optimal direction of migration. Interestingly, these extensive protrusions were completely absent in SEPT7 cKO cells, indicating that these cells lost their ability to properly form actin-based lamellipodia. In addition, we also observed fewer intercellular actin-based protrusions in 2D-adherent SEPT7-depleted cells as compared to control cells (Fig. S8). These observations suggest that septins stimulate the formation of lamellipodia in breast cancer cells.

A similar observation was previously made in MCF-7 breast cancer cells, where SEPT9 depletion inhibited protrusion formation, while overexpression of the SEPT9i1 isoform promoted the formation of lamellipodia and filopodia^36^. In endothelial cells and fibroblasts, septins have also been implicated in lamellipodia formation^49, 75^. The mechanisms are not well understood. It has been suggested that septins support membrane protrusions by preferential binding to regions of positive membrane curvature^75^. Also, reconstituted SEPT2/6/7 septin hexamers were recently shown to promote actin polymerization through septin-bound Cdc42EP3, a Cdc42 effector protein (also known as BORG2)^16^. Although we did not find any change in Cdc42 expression in Hs578T cells upon SEPT7 depletion by Western Blot analysis (Fig. S10C), it would be interesting to study the potential involvement of Cdc42 and BORG proteins. In A7 human melanoma cells, SEPT2 was recently shown to suppress lamellipodia formation through ARHGAP25, a GTPase-activating protein for the Rho small GTPase^56^. Moreover, SEPT2 and SEPT9 were shown to form a complex with the Rho-GTPase activating protein ARHGAP4 that regulate microenvironment-dependent focal adhesion reorganizations^76^. Thus, the impact of septins on protrusion formation likely depends on the composition of septin hetero-oligomers in the cell and on the cellular context. More in-depth analysis of septins and Rho and Rac signaling in Hs578T cells (and cancer cells more generally) will be needed to dissect this.

Since septins have been reported to promote cancer cell invasion through matrix metalloprotease secretion^38^, we also investigated cell shape control in rigid micropatterned environments where the cells are unable to actively remodel their environment. On micropatterned quasi-1D lanes that mimic cell adhesion along collagen fibers, we observed that SEPT7 depletion led to reduced cell aspect ratios. In microfluidic pillar devices that mimic the 3D pore space of collagen gels, the SEPT7-depleted cells were also less elongated than control cells. Moreover, they migrated with rounder and less branched cell shapes through the constrictions, similar to the cell behavior in collagen gels. However, overview images of microfluidic constriction areas qualitatively did not show clear differences in the number of migrating cells for control and SEPT7-depleted conditions (Fig. S6). The inhibitory effects of SEPT7 depletion on single-cell invasion of collagen networks that we observed may therefore involve a contribution of changed matrix remodeling besides protrusion regulation.

Although SEPT7 cKO cells had different shapes in collagen gels as compared to control cells, confocal reflection imaging of the collagen network indicated that the cells still interacted and actively pulled on the collagen fibers. Western Blot analysis showed that SEPT7 depletion indeed did not alter the expression of the collagen-binding integrin-β1 receptor (see Fig. S12A(iv)), which was confirmed by immunocytochemistry (see Fig. S12B). Likewise, expression levels of focal adhesion kinase, paxillin and vinculin were identical between control and SEPT7 cKO cells (see Fig. S12A(i-iii,v-vi,viii)). However, the ratio of phospho-paxillin/paxillin was increased about 1.5-fold in SEPT7 cKO cells compared to NT CTRL cells (see Fig. S12A(vii)). It will be interesting to study the relation between septins and focal adhesions in 3D-confined cells more closely in future, especially in light of recent evidence for interactions of paxillin with SEPT9i1 in 2D adhererent MCF-7 breast epithelial cells^36^ and with SEPT7 in U2OS cells and fibroblasts^49, 77^. In several cellular contexts, septins have been reported to affect the epithelial-to-mesenchymal (EMT) status of the cells^38, 78^. Western blot analysis of the Hs578t cells did not show any significant changes in the levels of key EMT markers (vimentin and N-cadherin) upon SEPT7 depletion (Fig. S10D, G).

Collectively, our findings show that septins regulate confined invasion of breast cancer cells by controlling cell shape and supporting actin-based protrusion formation to explore the extracellular matrix. The strong impact of septins on cell shape is interesting in light of evidence that cell shape indicators for complexity and protrusions correlate with triple negative breast cancer tumor growth and metastasis in *in vivo* and *in vitro* models^79, 80^. We still understand little about the molecular processes that govern cancer cell shapes and protrusive behavior, and how these are impacted by the biophysical properties of the tumor microenvironment. Our findings indicate an important role for a septin-mediated mechanism of cell shape and actin-based protrusion regulation that promotes 3D-tissue invasion.

## Resource availability

- Lead contact: Requests for further information and resources should be directed to and will be fulfilled by the lead contact, Gijsje Koenderink.
- Materials availability: All unique/stable reagents generated in this study are available from the lead contact without restriction.
- Data and code availability:

**–** All data reported in this paper will be shared by the lead contact upon request.
**–** Accession numbers are listed in the key resources table.
**–** Any additional information required to reanalyze the data reported in this paper is available from the lead contact upon request.

## Limitations of study

In this study we show the role of septin in breast cancer cell invasion using *in vitro* assays using 3D collagen gels and microfluidic constrictions to mimic the pore space that cells encounter in interstitial tissue. We note that our study was restricted to Hs578T breast cancer cells and to extracellular microenvironments that are structurally and biochemically much simpler than tissues. As cancer cell migration is a dynamic process with reciprocal cell-matrix crosstalk and interactions with stromal and immune cells, additional work using more complex *in vitro* and *in vivo* models is needed to understand the role of septin in 3D cancer cell invasion. Moreover, the molecular basis of the impact of septins on protrusion formation that we identified remains unclear. To dissect this, a more complete molecular characterization (for instance through proteomics and transcriptomics) is needed.

## Acknowledgements

A.vd.N., K.B., E.H.J.D. and G.H.K. gratefully acknowledge funding from the OCENW.GROOT.20t9.O22 project *The Active Matter Physics of Collective Metastasis* financed by the Dutch Research Council (NWO). M.T., R.C.B. and P.E.B. gratefully acknowledge funding from the European Research Council (ERC) under the European Union’s Horizon 2020 research and innovation program (Grant agreement no. 819424).

## Author contributions statement

AvdN: conceptualization, methodology, validation, formal analysis, investigation, data curation, writing – original draft, writing – review and editing, visualization. K.B.: methodology, validation, formal analysis, investigation, writing - review and editing. N.v.V.: investigation, validation, formal analysis, M.T.: investigation, formal analysis, writing - review and editing, M.B.: investigation, formal analysis, visualization, writing - review and editing, R.C.B.: conceptualization, methodology, writing - review and editing, P.E.B.: conceptualization, methodology, resources, writing - review and editing, E.H.J.D.: conceptualization, methodology, resources, writing - review and editing, project administration and funding acquisition, supervision, G.H.K.: conceptualization, methodology, validation, resources, writing – review and editing, project administration and funding acquisition, supervision.

## STAR Methods

### Experimental model and study participant details

#### Cell culture and generation of inducible SEPT7 knockout cells

Human breast cancer cells (Hs578T, #HTB-126, ATCC) were cultured in Dulbecco’s Modified Eagle Medium (DMEM, Thermo Fisher, #11574486) supplemented with 5% Fetal Bovine Serum (Thermo Fisher Scientific, #11573397) and 1% penicillin-streptomycin (Thermo Fisher Scientific, #11528876). Cells were subcultured between 80-90% confluency and incubated at 37°C and 5% CO_2_. Cells were periodically checked for mycoplasm.

A doxycycline-inducible Hs578T Cas9 cell line was produced using Edit-R Inducible Lentiviral hEF1α-Blast-Cas9 Nuclease Plasmid DNA (Dharmacon, CAS11229, CO, USA). A subconfluent monolayer of Lenti-X 293T cells (Clontech, #632180, CA, USA) was adhered to a 10 cm dish overnight. A DNA mixture was prepared with third generation lentiviral helper vectors using 2.6 µg pMDLg-RRE, 1.4 µg pCMV-VSVG and 2.0 µg pRSV-Rev (all from Addgene, #12251, #8454, #12253, MA, USA) and 4.0 µg of the Cas9 plasmid. Lenti-X 293T cells were transfected with the DNA mixture and 0.05 mg Polyethylenimine (Polysciences, #23966-2, PA, USA) in 10 mL culture medium. After 24 hours the medium was refreshed, after 48 hours and 72 hours the medium was collected and filtered through a 0.45 µm filter. 50,000 Hs578T cells were adhered in a 6-wells plate (Greiner Bio-one, #657160, Austria) and virus-containing medium was added and supplemented with 5 µg/mL Polybrene (Sigma-Aldrich, MO, USA). After 24 hours the medium was changed and cells were selected using 2 µg/mL Blasticidin (Sigma-Aldrich, #203350, MO, USA). A single clone of Hs578T-Cas9 was isolated, expanded and tested for Cas9 expression upon Doxycycline (Selleckchem, #S5159, TX, USA) exposure. For sgRNA plasmids, the following guide RNA was added into the U6-gRNA/PGK-Puro-2A-BFP vector (Addgene, #50946): SEPT7: ’CCTGTTATCGACTACATTGATAG’; Additionally a non-targeting sgRNA was used as negative control. HEK293T cells were transfected with the pKLV-U6-gRNA/PGK-Puro-2A-BFP vectors cloned with sgSEPT7 or the non-targeting control sgRNA, with 2.5 mg/mL polyethylenimine (Polysciences, #23966-2) in phosphate-buffered saline (PBS, Gibco). Lentivirus-containing medium was collected from the HEK293T cells for 2 days and used to transduce Hs578T-Cas9 cells. Selection of transduced cells was performed with 1 µL puromycin (Thermo Fisher, #A1113803) for two days.

For induction of the CRISPR-Cas9 knockout, media was supplemented with 1 µg/mL doxycycline (Bio-connect, S5159), creating Hs578T cells with a SEPT7 knockout (Hs578T SEPT7 cKO) and a non-targeting control (Hs578T NT CTRL). We found through immunofluorescence (Fig. S1) and Western Blot analysis (Fig. S2) that a minimum of 7 days was needed for maximal SEPT7 depletion. Knockout cell cultures were used for a maximum of three weeks to prevent the influence of long-term effects of the knockout and to prevent overgrowth of remaining septin-positive cells in the culture.

### Method details

#### Actin polymerization inhibition

In experiments requiring actin depolymerization, we incubated the cells for 30 minutes with culture media supplemented with Cytochalasin D (Merck Sigma, #C2618) or with 0.025 µL/mL DMSO (Bioke, #12611P) as a vehicle control. This drug concentration is sufficient to cause disappearance of F-actin stress fibers in Hs578T cells^81^.

#### Microfluidic migration assays

To probe the impact of SEPT7 knockout on cell shapes during confined migration, we employed our recently established polydimethylsiloxane (PDMS)-based microfluidic device with custom-designed constriction areas^82^, which was adapted from earlier work of Davidson et al.^83^. The multi-layered master mold was created using standard soft lithography at the Kavli Nanolab Delft, with a µMLA laserwriter (Heidelberg Instruments). The design consists of a 5 µm tall and 440 µm wide constriction area aligned with an adjacent 50 µm tall perfusion channel for cell loading, chambers that end at the constriction area, and a bypass channel to equilibrate the fluid levels between the reservoirs positioned at the outer sides of the chamber.

The first layer of the design consisted of 5 µm SU-8 2005 photoresist (Kayaku Advanced Materials) and was spun on a 4-inch silicon wafer that was soft-baked and post-baked at 95 °C for 2 minutes. The second layer consisted of 45 µm SU-8 3050 (Kayaku Advanced Materials) and was soft-baked at 95 °C for 15 minutes, post-baked for 1 minute at 65 °C and 5 minutes at 95 °C, and developed with a SU-8 developer (Sigma Aldrich). After developing, trichloro(1H,1H,2H,2H-perfluoroctyl)silane (Sigma Aldrich) was coated on the master mold. Microfluidic chips were made from PDMS (Sylgard 184, Dow Corning), prepared with a curing agent with a 10:1 (w/w) ratio. The PDMS was poured on the silicon mold, degassed and cured at 65 °C for 3 hours. Reservoirs and cell-loading ports were punched with a revolving punch plier (Knipex) and a 0.75 mm diameter punch (Rapid-core, Welltech), respectively. PDMS chips and glass coverslips were plasma cleaned (Harrick Plasma) at 30 W for 150 seconds and bonded overnight at 65 °C.

For live-cell experiments, microfluidic chips were sterilized with 70% ethanol and subsequently washed three times with MQ and once with PBS. Chips were coated with collagen by adding 100 µL 100 µg/mL PureCol type 1 bovine collagen (Advanced Biomatrix) in PBS through the cell-loading ports and incubating for 2 hours at room temperature. Chips were washed thrice with PBS and once with cell culture medium. Next, a suspension of trypsinized cells (6 µL containing 30,000 cells) was added to each chip via the cell-loading ports. The reservoirs were filled with cell culture media and the chips were incubated overnight in a cell culture dish at 37°C and 5% CO_2_, together with a 15 mL falcon tube cap filled with MQ to prevent evaporation. One hour before imaging, cell culture medium was removed from the reservoirs and replaced with cell culture medium containing live-cell dyes to stain the nucleus (Hoechst 33342, Thermo Fisher, 1:10,000 solution) and the cytoplasm (Cytotracker Orange, Thermo Fisher, 1:1,000 dilution). After one hour incubation, the device was sealed with a glass coverslip and imaged by confocal microscopy.

### Micropatterning

For micropattern fabrication, we adapted a well-established deep UV printing protocol^84^. Coverslips were first cleaned with isopropanol (VWR Chemicals, #20922.264) and Milli-Q (MQ) and sonicated for 5 minutes in an Ultrasonic Cleaner (Emerson, Brandson 2510), followed by 5 minutes in a UV plasma cleaner (BioForce Nanoscience, UV Ozone ProCleaner Plus). PLL-PEG coatings were made using 1 mgmL^−1^ PLL-g-PEG (Poly(L-lysine)-graft-poly(ethylene glycol), Susos AG) diluted 1:10 in 10 mM HEPES buffer, pH 7.4 (Thermo Fisher, #11560496) and pipetted in 50 µL droplets on parafilm. Cleaned coverslips were placed on top of the droplets and incubated at room temperature for 1-2 hours. Next, coated coverslips were washed 10x in MQ and dried using a nitrogen spray gun.

The photomask was fabricated by DeltaMask BV. The photomask was first cleaned with acetone and isopropanol and next plasma cleaned for 10 minutes in the UV plasma cleaner. To attach the coverslips to the photomask, 1.5 µL droplets were pipetted onto the patterns before adding the coverslips on top. The patterns were printed in the UV plasma cleaner for 10 minutes. Patterned coverslips were sterilized with 70% ethanol and washed twice with PBS. Patterned coverslips were incubated with 100 µL 10 µg/mL Rhodamine-labeled fibronectin (Universal Biologicals, #FNRO1-A) in PBS for 1 hour. Next, coverslips were washed with PBS and stored in PBS at 4 °C for a maximum of 1 week. To seed the cells on the patterns, patterned coverslips were placed in 6-well plates and seeded with a density of 50,000 cells/well for at least 24 hours to ensure attachment.

### Spheroid invasion in collagen gels

Bovine hide collagen type I (purity ≥ 99.9%, Advanced Biomatrix, provided as a solution of 3 mg/ml collagen in 0.01 N HCL) was used to prepare 2.4 mg/ml collagen gels. The gels were made isotonic by adding 12.5 v/v% of 10x Phosphate Buffered Saline (PBS, Thermo Fisher). Next, 0.1 M sodium hydroxide was added to bring the pH to 7.4. The solution was further diluted with pre-cooled MQ and vortexed for 30 seconds and polymerized at 37°C in a µ-slide 8-well (Ibidi) for 45 minutes.

For spheroid formation, cells were seeded in round-bottom and ultra-low attachment Elplasia*^TM^* 96-well plates (Corning), with a seeding density of 40,000 cells/well to create spheroids with an average size of ∼200 µm. Spheroids were incubated for 2-3 days to allow spheroid formation and compaction, with a small necrotic core, and pipetted out of the wells using a cut-off 200 µL pipet tip to prevent shearing the spheroids. Spheroids were pipetted on top of 80 µL pre-polymerized collagen in each Ibidi well and incubated for 45 minutes. Medium was pipetted out of the well, and another 100 µL collagen was polymerized on top of the spheroids, creating a sandwich model, adapted from Ref.^85^. The collagen samples were supplemented with culture medium containing Hoechst 33342 (1:10,000) and Cytotracker Orange (1:5000) in order to fluorescently tag the nucleus and cytoplasm of the cells, respectively.

### Immunocytochemistry

For immunocytochemistry analysis, Hs578t NT CTRL and SEPT7 cKO cells were seeded with a density of 200,000 cells/well in 6-well plates with 2 coverslips per well for 24-48 hours. For fixation of cells cultured on 2D substrates (coverslips or fibronectin-coated micropatterns), culture medium was removed and replaced with 4% para-formaldehyde (PFA, Tebu-Bio) in PBS. After a 10 minutes incubation, PFA was removed and the samples were washed three times with PBS. Next, cells were permeabilized with 0.5% Triton X-100 (Merck Sigma) in PBS for 3 minutes and washed three times with PBS supplemented with 0.1% Tween (PBS-T, Merck Sigma). The samples were blocked for 30 minutes with 3% Bovine Serum Albumin (BSA, Merck Sigma) in PBS.

Primary stainings were made in 1.5% BSA/PBS and incubated between 6-24 hours at 4 °C. Primary antibodies used were anti-α-tubulin (1:1000, Invitrogen, #32-2500), anti-gm130 mouse (1:500, BD Biosciences, #610823), anti-septin-7 rabbit IgG (1:250, Thermo Fisher, #PA5-56181) and anti-septin-9 rabbit IgG (PAN-septin-9, 1:200, Proteintech, #10769-1-AP). After primary staining, samples were washed three times with PBS-T.

Secondary stainings were made in PBS-T and incubated between 1-6 hours at room temperature. Secondary antibodies and dyes used were anti-rabbit goat IgG 488 (1:1000 Thermo Fisher, #A11008), anti-mouse goat IgG 568 (1:1000, Thermo Fisher, #a11004), anti-rabbit goat IgG 647 (1:1000, Thermo Fisher, #a27040), Hoechst 33343 (1:1000) and Phalloidin 647 (1:250). After secondary staining, samples were washed three times with PBS-T and once with MQ. Coverslip samples were mounted on microscope slides (Thermo Fisher, #16309475) using ProLong Diamond mounting solution (Thermo Fisher, #15468070), air-dried and stored at 4 °C.

Spheroid-collagen samples were stained with the same protocol as described above, but with increased incubation times and without mounting the samples. Instead, the samples were stored in PBS at 4 °C for a maximum of two weeks. Incubation times were adapted for BSA blocking (overnight), primary staining (overnight), secondary staining (overnight) and wash steps (30 minutes). Samples were stored in PBS at 4 °C.

### Microscopy imaging and automated image analysis

Confocal fluorescence and reflectance imaging was done on a Stellaris 8 confocal microscope (Leica), equipped with a supercontinuum white light laser, 405 nm laser and three hybrid detectors. Immunocytochemistry samples were imaged with 40x/1.25 or 63x/1.30 glycerol objectives and the 405 nm laser and 488/553/650 nm laser lines for fluorescence excitation. Cells on glass (2D samples) were imaged ∼ 1 µm above the glass and cells in collagen (3D samples) were imaged with 0.25 µm Z-steps from top to bottom (Z-stack size ranging from 8.5 - 52.5 µm. Spheroid samples were imaged around the spheroid equator with a Z-stack size of 70 µm and Z-step size of 10 µm. Live-cell imaging of microfluidic chips and spheroid assays was performed with a 20x/0.75 air objective and the 405 nm laser and the 458 nm laser line for fluorescence imaging of the cells and 488 laser line for reflection imaging of collagen. Time-lapse image series at multiple locations were acquired with time intervals of 15 minutes over a total period of 24-72 hours. Environmental control regulated the temperature at a constant 37 °C and 5% CO_2_. Spheroids in collagen were imaged at the spheroid equator and microfluidic chips were imaged ∼ 1 µm above the glass.

Automated image analysis to detect 2D cell shape parameters was done in in Fiji^86^ in which we binarized Cytotracker Orange (for microfluidic time-lapses) and actin signals (for immunofluorescence images) from which we measured cell area, aspect ratio, circularity and solidity. To measure lengths of cells on 2D substrates, we analyzed the skeleton with the ’skeletonize’ plugin in Fiji and measured the longest path of the skeleton. In 3D samples, cells had a variety of orientations, complicating quantitative analysis of cell lengths and shapes. We therefore used the ’AnalyzeSkeleton’ plugin of Fiji^86, 87^ on binarized maximum projections of actin intensities. From the skeleton, we could count the number of junctions in the skeleton as a measure for protrusions, and extract the longest path in the skeleton to determine the cell length in the XY plane. To also determine the length in Z, we counted the number of Z-stacks in which the cell was imaged. Cell elongation was defined as the combination of measured cell lengths in XY and in Z. Cells were selected when located a vertical distance > 20 µm from the coverglass. Maximum time projections of time-lapse imaging series of 3D spheroid invasion were obtained from maximum projections of the colour inverted images of the time-lapses in Fiji^86^.

### Western Blot analysis

For the Western Blot analysis of the SEPT7 cKO induction with doxycyclin treatment, Hs578T NT CTRL and SEPT7 cKO cells were seeded at a cell density of 300,000/well in 6-well culture plates. After a minimum of 24 hours, cells were washed with PBS and subsequently lysed in radioimmunoprecipitation buffer (RIPA, 100 µL/well, Thermo Fisher) and stored at -20°C. Loading samples made with Laemmli buffer (2x, Bio-rad) and 4% β-mercaptoethanol (Sigma Aldrich) were boiled at 95°C for 5 minutes. Mini-PROTEAN-TGX gels (Bio-rad) were used for sodium dodecyl sulfate-polyacrylamide gel electrophoresis (SDS-PAGE) at 100 V for 1.5 hours. Western Blotting of the gels was done on Trans-Blot Turbo Mini 0.2 µm PVDF Transfer Packs (Bio-rad) using a Trans-Blot Turbo Transfer System (Bio-rad). After blotting, membranes were blocked in 5% BSA/PBS overnight. Membranes were stained with primary antibodies: anti-GAPDH rabbit IgG (1:1000, #CST2118S, Bioke), anti-septin-2 rabbit IgG (1:1000, Sigma, #HPA018481), anti-septin-7 Rabbit IgG (1:1000, #18991, IBL-America) and anti-septin-9 rabbit IgG (PAN-septin-9, 1:10000, Proteintech, #10769-1-AP) in 5% BSA/PBS overnight, on a shaker at 4°C. After primary stainings, membranes were washed thrice with PBS-T. Secondary stainings were done with goat anti-rabbit HRP (#ab6728, Abcam), 1:5000 in PBS-T. Membranes were washed thrice with PBS-T. Imaging was executed with an enhanced luminol-based chemiluminescent substrate kit (Thermo Fisher) on a gel-imager (Bio-rad). Intensity bands were measured three times in Fiji^86^ and subtracted from different background spots.

For the Western Blot analysis of the SEPT7 cKO and NT CTRL Hs578T cell lines, both cell lines were seeded and grown with doxycyclin for 7 days. Cells were scraped in PBS and lysed in equal volumes of 2x Laemmli buffer (4% SDS, 20% glycerol, 120mM Tris pH 6.8) with Protease and Phosphatase Inhibitor Cocktail (PPC1010, Sigma). Lysates were cleared of large DNA by passing through a 25H needle and then heated to 65°C for 10 minutes. Protein concentrations were measured with the Lowry protein assay. Equal amounts of protein were size separated by SDS-PAGE containing tricloroethylene (TCE). After running, the gel was exposed to a 300 nm transilluminator (Bio-Rad ChemiDoc Imaging System) for 5 minutes. Western Blotting was done on Trans-Blot Turbo mini 0.2 µm PVDF Transfer Packs using a Trans-Blot Turbo Transfer System. The blots were imaged for TCE on the transilluminator. Membranes were then blocked for 1 hour with 5% BSA in PBS supplemented with 0.1% Tween-20. Membranes were incubated overnight at 4°C with primary antibodies: anti-actin mouse IgG (1:50,000, #MAB1501, Millipore), anti-anillin mouse IgG1 (1:1000, #MA5-31342, Invitrogen), anti-CDC42 rabbit IgG (1:10,000, #AB187643, Abcam), anti-FAK mouse IgG1 (1:500, #610088, BD Biosciences), anti-GAPDH rabbit IgG (1:5000, #2118s, Cell Signaling), anti-integrin-β1 rabbit IgG (1:1000, #9699t, Abcam), anti-MLC2 rabbit IgG (1:1000, #8505T, Cell Signaling) anti-N-cadherin rabbit IgG (1:4000, #22018-1-AP, ProteinTech), anti-paxillin mouse IgG1 (1:200, #AHO0492, Invitrogen), anti-pMLC2 rabbit IgG (1:1000, #3671T, Cell Signaling), anti-phospho-paxillin rabbit IgG (1:500, #44-722G, Invitrogen), anti-septin-7 rabbit IgG (1:1000, #18991, IBL), anti-α-tubulin, mouse IgG (1:2000, #32-2500, Invitrogen), anti-vimentin mouse IgG1 (1:20,000, #AB8978, Abcam) and anti-vinculin rabbit IgG (1:500, #700062, Thermofisher). After primary stainings, membranes were washed 5 times with PBS-T and incubated for 1 hour with secondary antibodies in PBS-T: rabbit anti-mouse HRP (1:5000, #AB6728, Abcam) and goat anti-rabbit HRP (1:5000, #AB97051, Abcam). Imaging was executed with an enhanced luminol-based chemiluminescent substrate kit on a ChemiDoc Imaging System (Bio-rad).

### Dynamic compression of spheroids

To measure the viscoelastic properties of the NT CTRL and SEPT7 cKO spheroids, dynamic compression was imposed on spheroids using a microfluidic constriction device with a constriction width of 84 µm in length of 110 µm described in Ref.^73^. For spheroid formation, NT CTRL and SEPT7 cKO cells were seeded in round-bottom and ultra-low attachment Elplasia*^TM^* 96-well plates, with a seeding density of 60,000 cells/well. Spheroids were incubated for 3 days and pipetted out of the wells using a cut-off 200 µL pipet tip. Deformation of the spheroid in the constrictions was imaged on an inverted wide field fluorescence microscope (Zeiss Axio-Observer) with camera streaming using a 5xNA 0.16 air objective and Zeiss Axio-Observer 0.63x digital camera. Experiments with more than one spheroid present in the microfluidic device during compressions were not considered in the analysis. Automated image analysis of spheroid shapes was performed using a custom Matlab script that detects the spheroid boundary and centroid, to calculate velocity and axial strain (see Ref.^73^ for details). The viscoelastic properties were extracted from the axial strain (derived from the axial spheroid dimensions) evolution over time, which characterized the spheroid deformation, via the fitting of a Dynamic Modified Maxwell Model (DMMM). The DMMM was derived by defining an applied stress *σ* in the Modified Maxwell Model by pressure differences across the channel, accounting for time-dependent pressure differences present in the system (the detailed mathematical derivations are provided in previous work Ref.^73^).

### Quantification and statistical analysis

Statistical analysis was performed using Microsoft Excel. Two-tailed Student’s t-tests were performed using the TTEST function. Statistical details of experiments are found in the figure legends and method details. P-value results from t-tests are indicated by: (ns) = p≥0.05, (*) = p<0.05, (**) = p<0.01, (***) = p<0.001. Error bars represent the standard error of the mean.

## Supplementary Information

**Figure S1.**
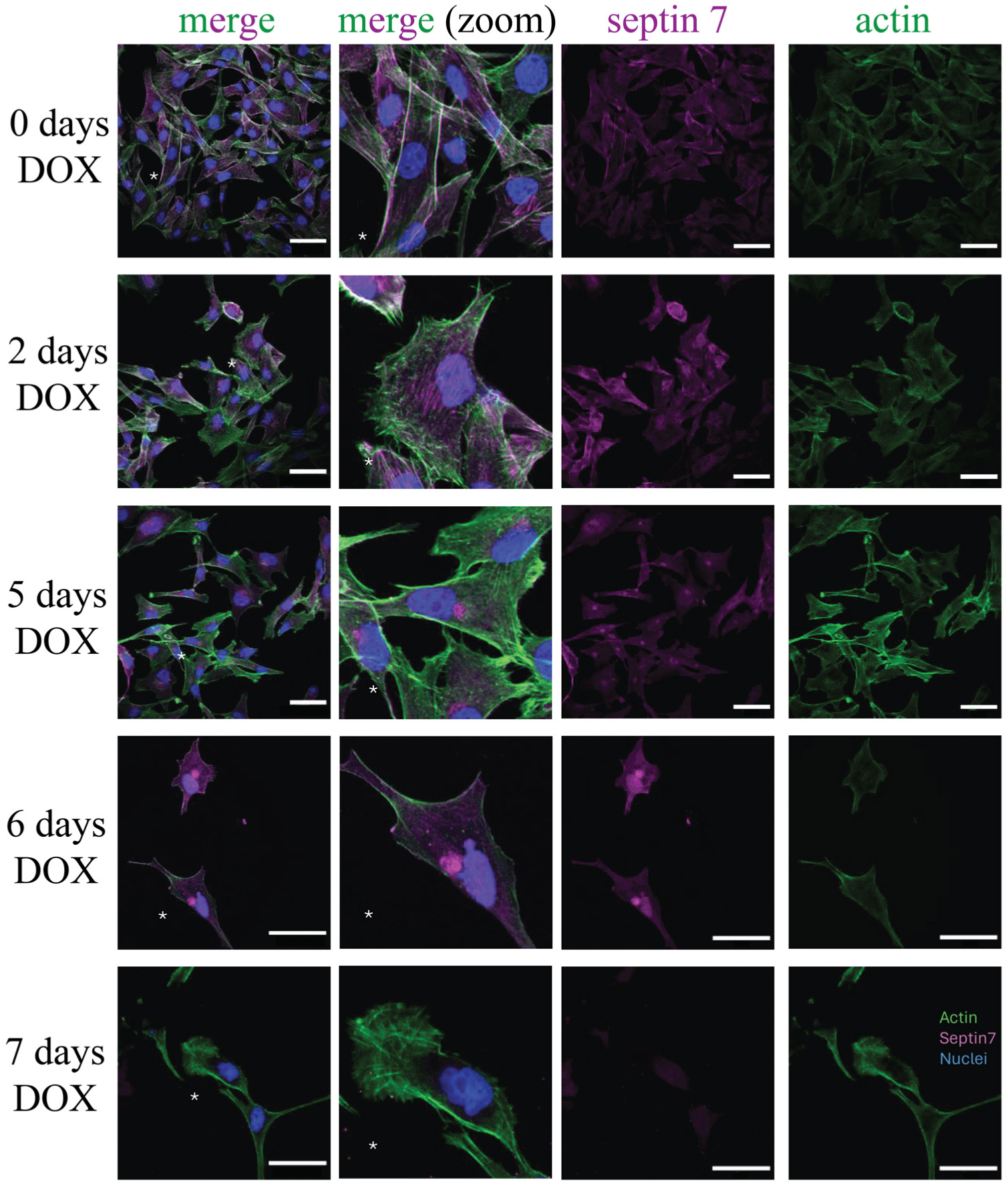
Immunocytochemistry analysis of the time dependence of doxycyclin (DOX) induction of SEPT7 knockout in Hs578T cells. Rows show cells after DOX induction over periods of (from top to bottom): 0, 2, 5, 6 and 7 days. Columns show stainings for actin (green), SEPT7 (magenta), merge of actin and SEPT7, and zoom-in of merge images at spots indicated with white asterisks. Nuclei are stained with Hoechst and displayed in blue. Image and display settings are the same for all images. Scale bars are 50 µm.

**Figure S2.**
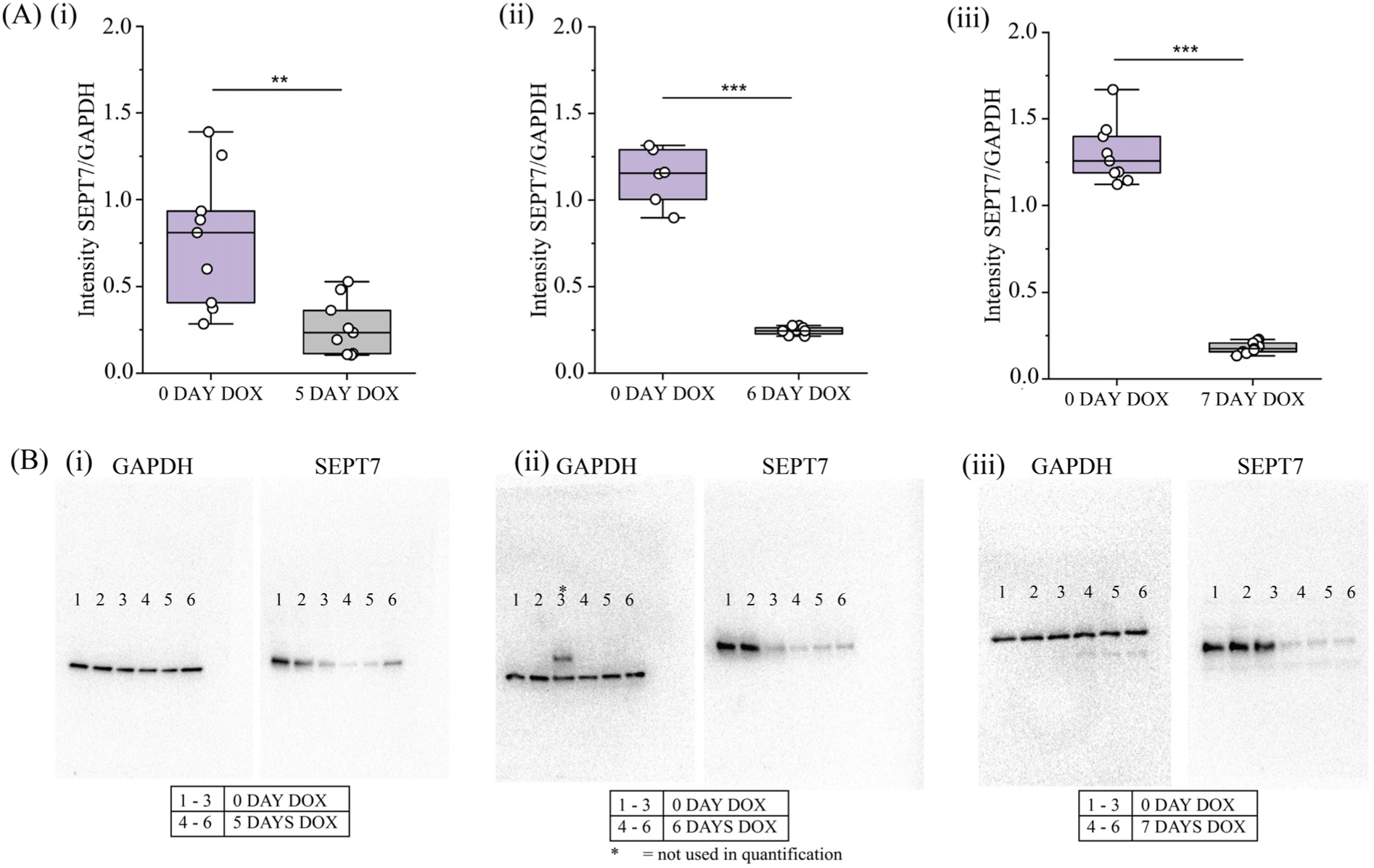
Western Blot analysis of the time dependence of doxycyclin induction of SEPT7 knockout in Hs578T cells. (A) Quantification of SEPT7 signal intensities normalized to GAPDH signal intensities for (i) 0 and 5 days doxycyclin treatment, (ii) 0 and 6 days doxycyclin treatment and (iii) 0 and 7 days doxycyclin treatment. (**) = p < 0.005, (***) = p < 0.001. (B) Western Blots showing GAPDH and SEPT7 stainings for (i) 0 and 5 days doxycyclin treatment, (ii) 0 and 6 days doxycyclin treatment and (iii) 0 and 7 days doxycyclin treatment. Note that sample 3 on panel B(ii) is not used for quantification because of the double GAPDH band.

**Figure S3.**
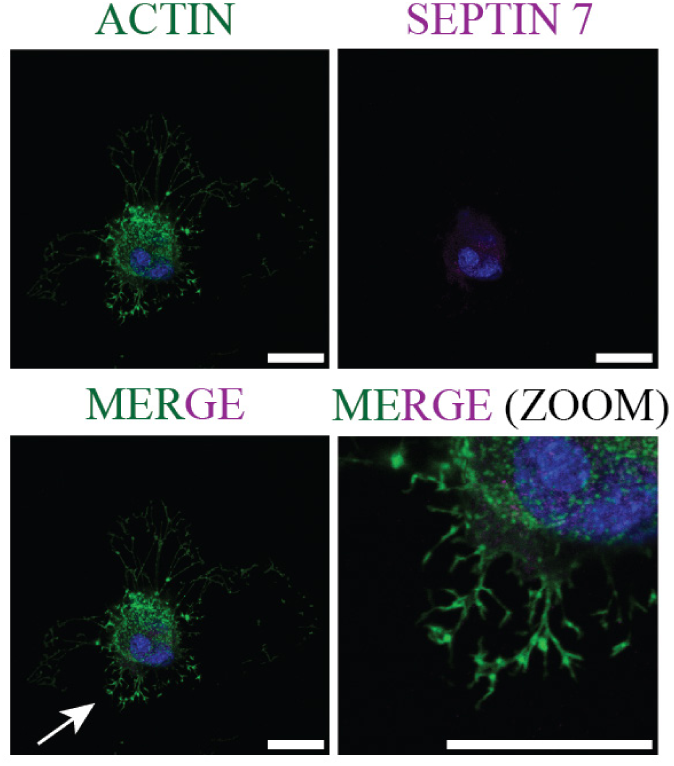
Inhibition of actin polymerization with cytochalasin D combined with SEPT7 depletion in Hs578t cells results in anomalous cell shapes with small 2D-adhered cell sizes. Immunocytochemistry images of SEPT7 cKO cells treated with cytochalasin D for actin (green) and SEPT7 (magenta). White arrow in MERGE image indicates area used for MERGE (ZOOM) images. Quantification of the reduction in cell area upon SEPT7 depletion is shown in Fig. 2. Scale bars are 25 µm.

**Figure S4.**
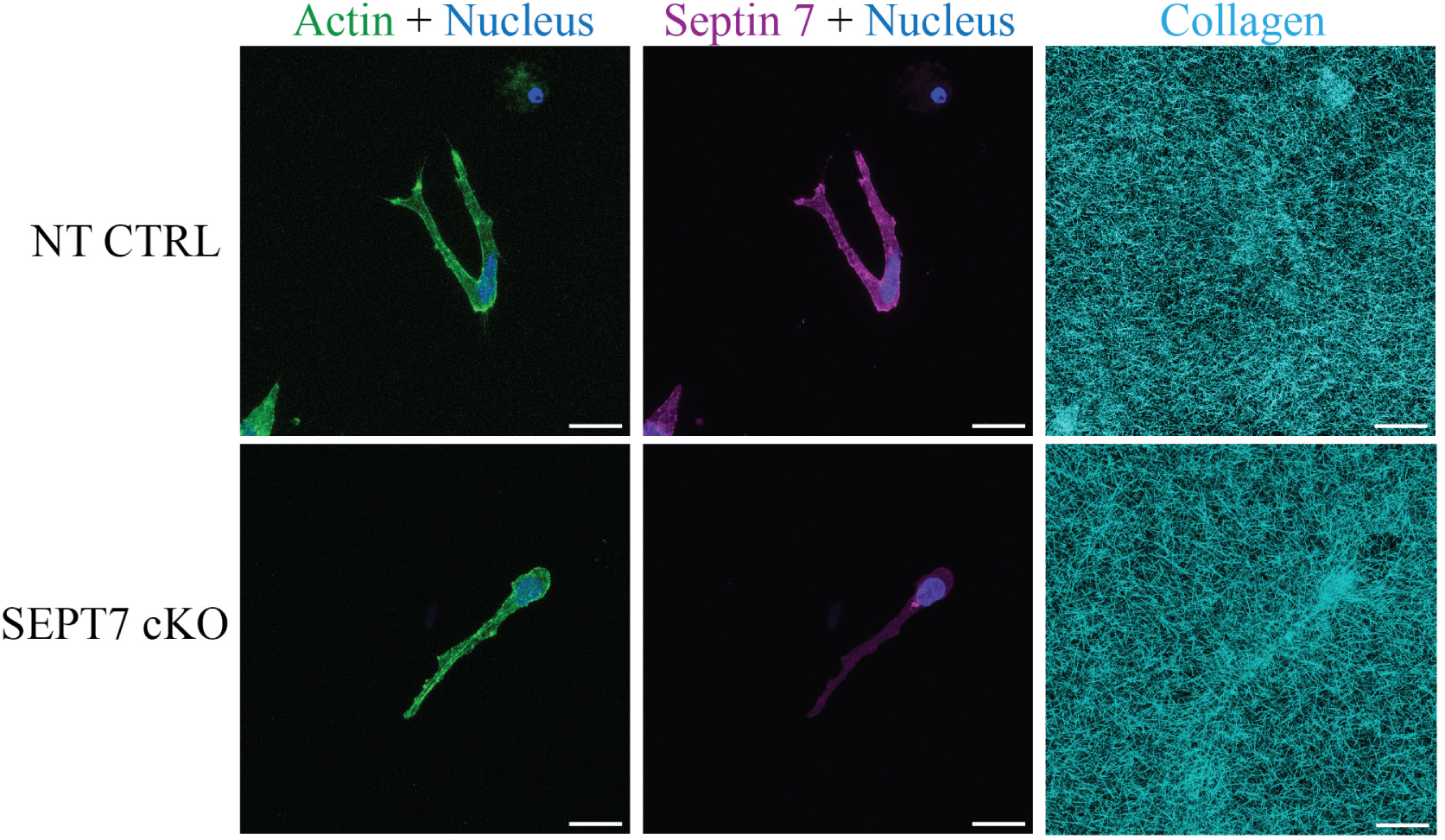
Images of Hs578T cells embedded in collagen network. Maximum intensity Z-projections of confocal fluorescence images of NT CTRL and SEPT7 cKO cells in a 2.4 mg/mL collagen gel after 48 hours. Cells were stained for nuclei (blue), actin (green) and SEPT7 (magenta). Collagen (blue) was imaged by reflection microscopy. Scale bars are 25 µm.

**Figure S5.**
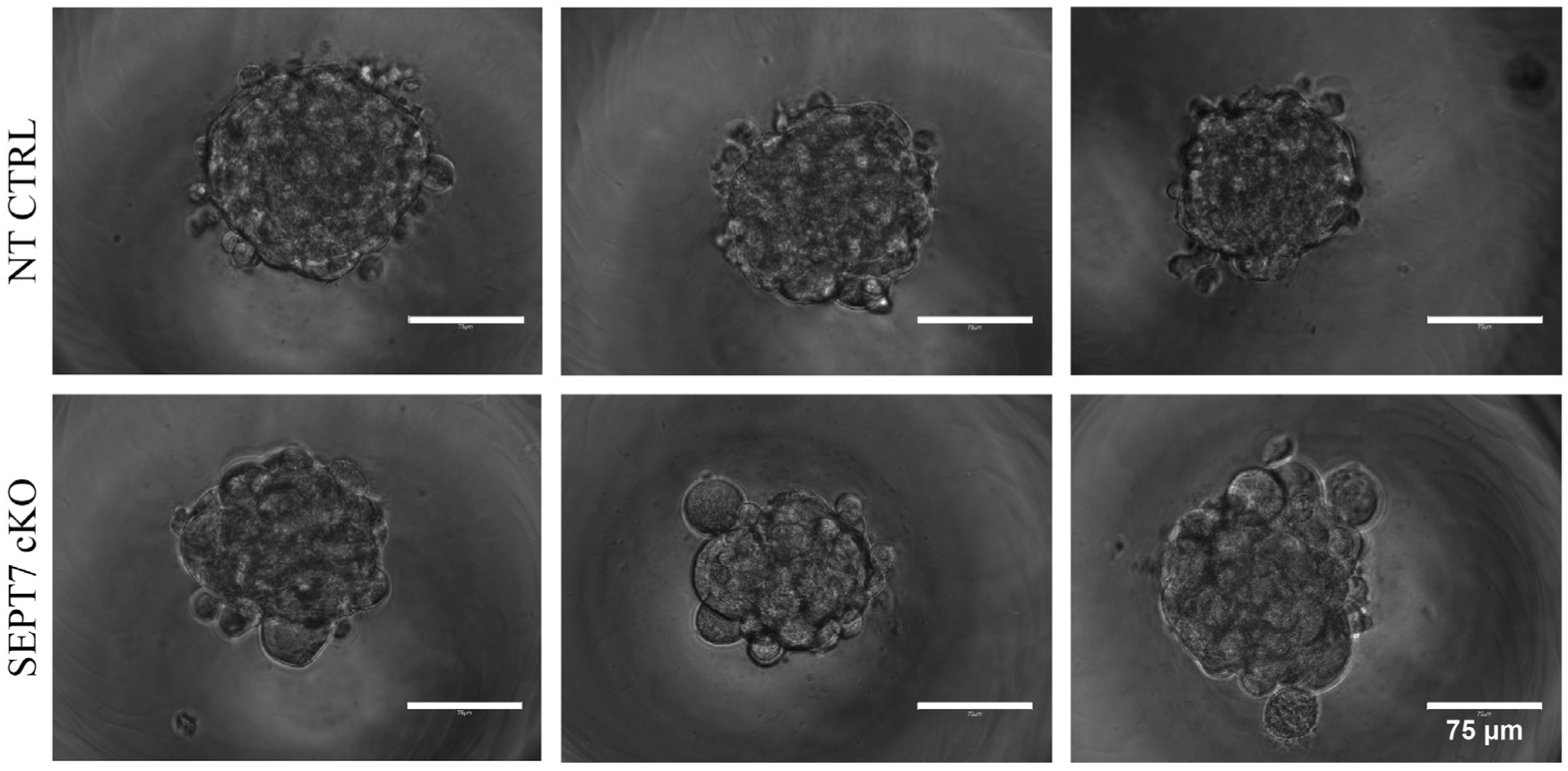
Morphological characterization of Hs578T cell spheroids. Brightfield images of NT CTRL (top) and SEPT7 cKO (bottom) Hs578T spheroids formed in round-bottom and ultra-low attachment Elplasia*^TM^* 96-well plates after 48 hours of incubation. Three representative examples are shown in each case. Note that the SEPT7 cKO spheroids are less round and exhibit increased edge roughness compared to NT CTRL spheroids. Scale bars are 75 µm.

**Figure S6.**
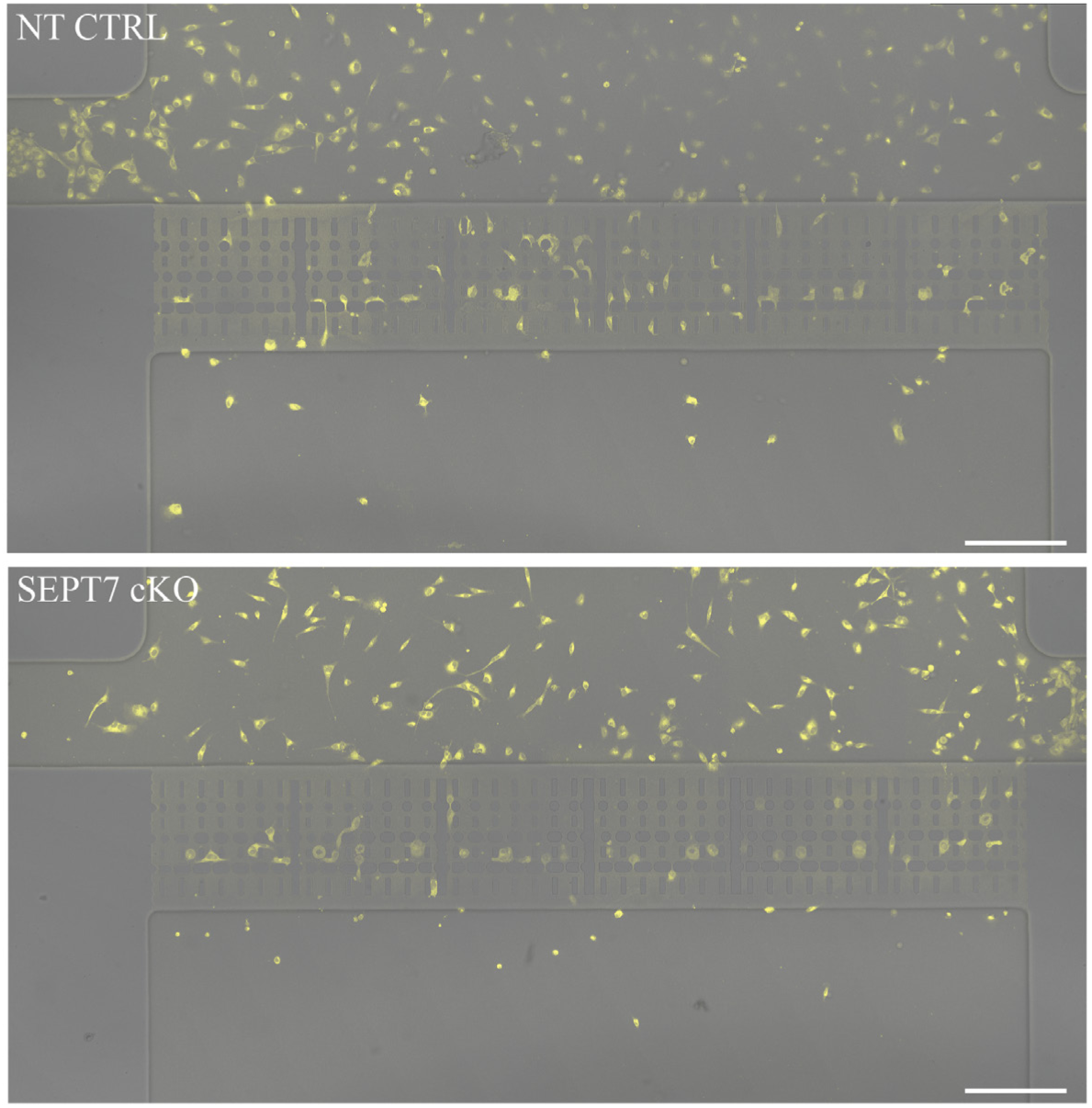
Overview images of cells in microfluidic migration chips. Brightfield images of microfluidic migration chips for NT CTRL (top) and SEPT7 cKO (bottom) Hs578T cells after 24 hour incubation, merged with Cytotracker Orange (yellow) signals for cell labeling. Note that the number of cells within the constriction areas is comparable for NT CTRL and SEPT7 cKO cells. Scale bars are 50 µM.

**Figure S7.**
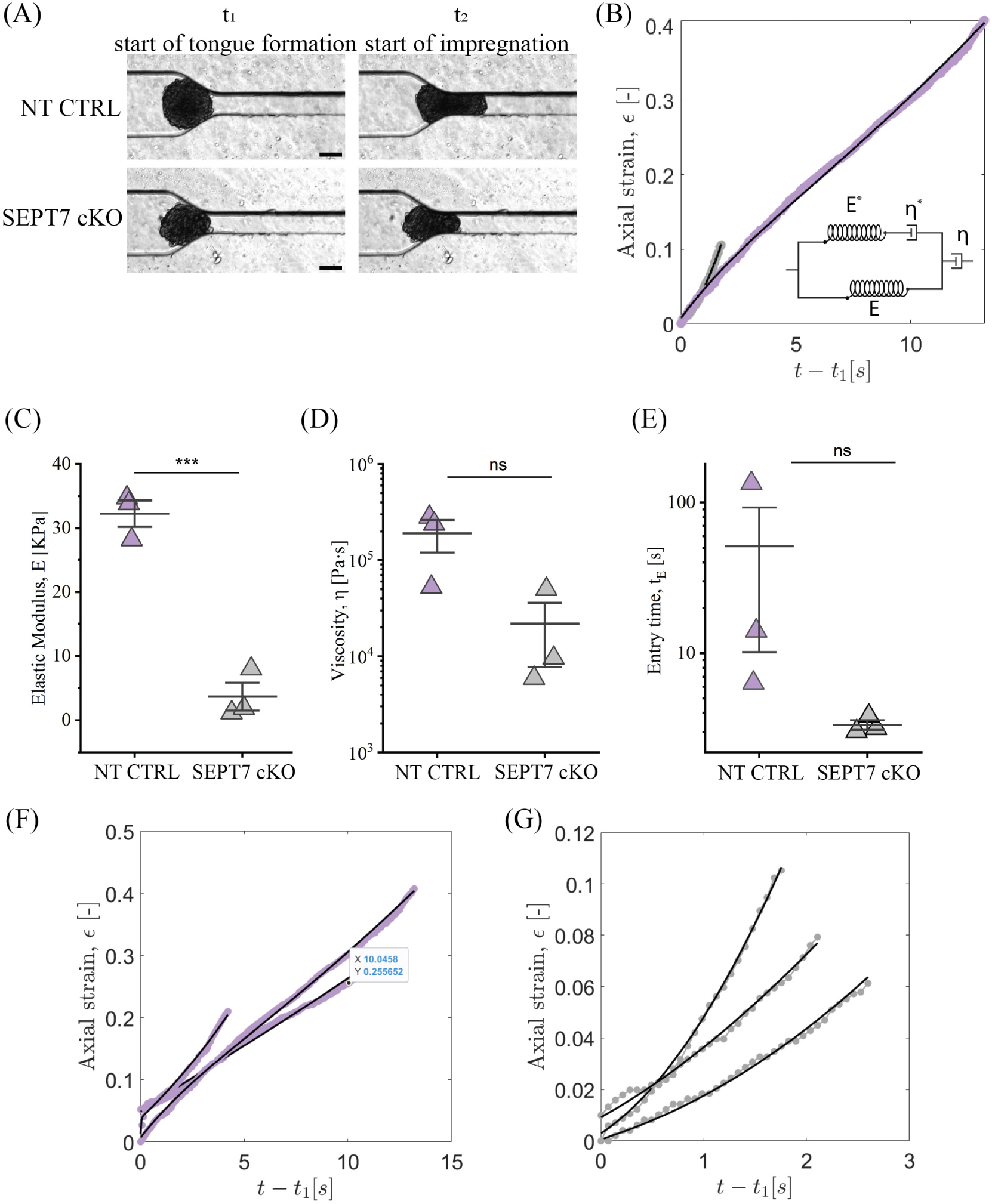
Microfluidic measurements of the viscoelasticity of Hs578T cell spheroids. (A) Brightfield images of NT CTRL and SEPT7 cKO Hs578T spheroids in the microfluidic constriction channel (for details of the device, see Ref.^73^): snapshots at time *t*_1_ (start of tongue formation) and *t*_2_ (start of compression). Scale bars are 100 µm. (B) Strain curves from NT CTRL and SEPT7 cKO Hs578T spheroids shown in panel (A). Strain curves start at the beginning of the tongue formation (*t*_1_) and are fitted to the Dynamic Modified Maxwell Model^73^, which is schematically illustrated in the inset. (C) Measurements for spheroid bulk elastic moduli *E*, (D) viscosity *η* and (E) entry time *t_E_*. (***) = p < 0.001, (ns) = p > 0.05. Bar and whiskers show the mean and standard error. N = 3 per condition. (F) All individual strain curves from NT CTRL spheroid compressions and (E) SEPT7 cKO spheroid compressions.

**Figure S8.**
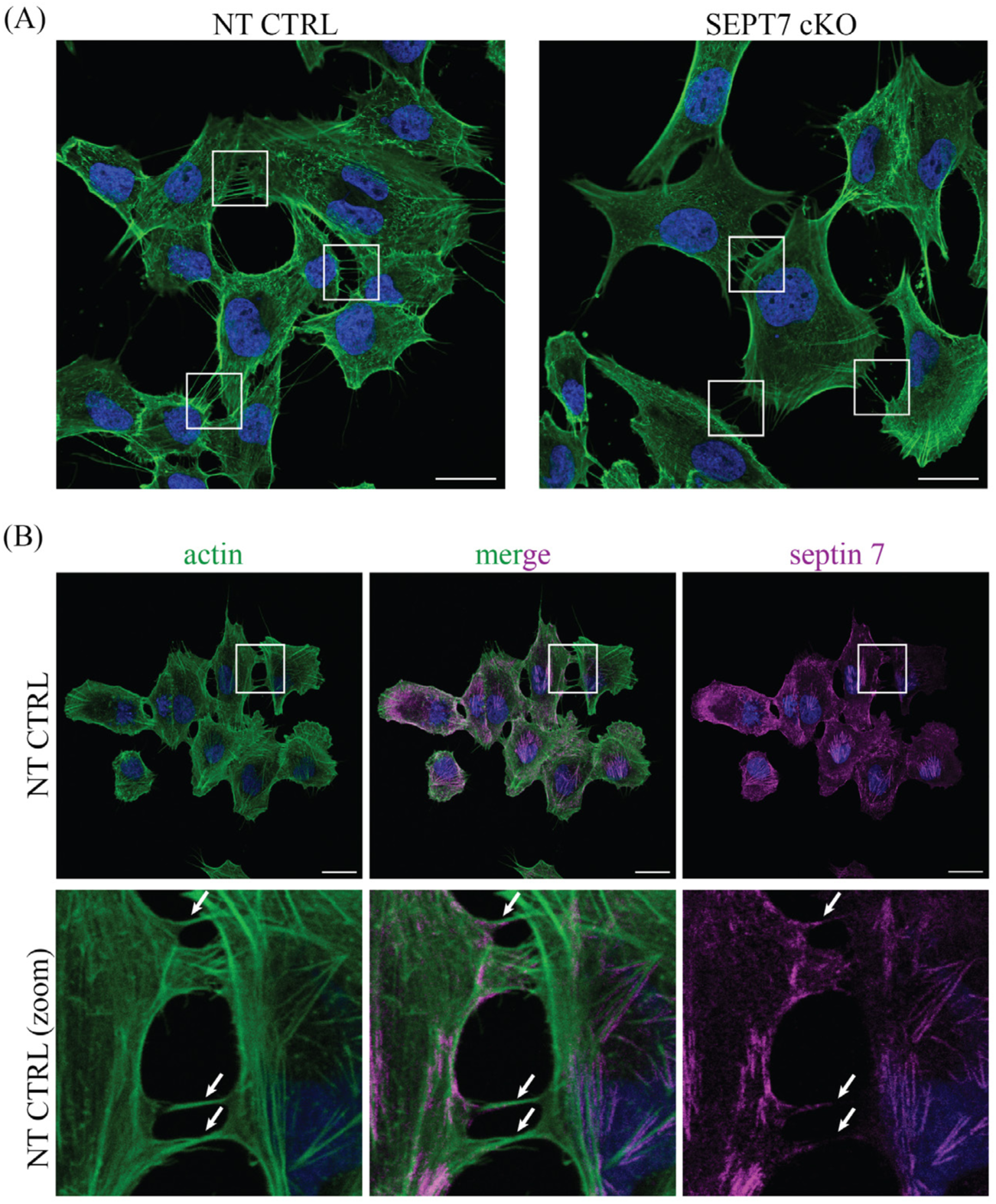
Intercellular actin-based protrusions in 2D-adherent Hs578T cells. (A) Immunocytochemistry of NT CTRL and SEPT7 cKO Hs578T cells stained for actin (green) and nuclei (blue). Connecting actin-rich protrusions between cells are reduced in SEPT7 cKO compared to NT CTRL, as seen in the areas indicated by white squares. (B) Immunocytochemistry of NT CTRL Hs578T cells stained for actin (green), SEPT7 (magenta) and nuclei (blue). Zoom in panels are indicated by white squares. White arrows indicate the connecting actin protrusions and co-localization with SEPT7 signals. Scale bars are 25 µm.

**Figure S9.**
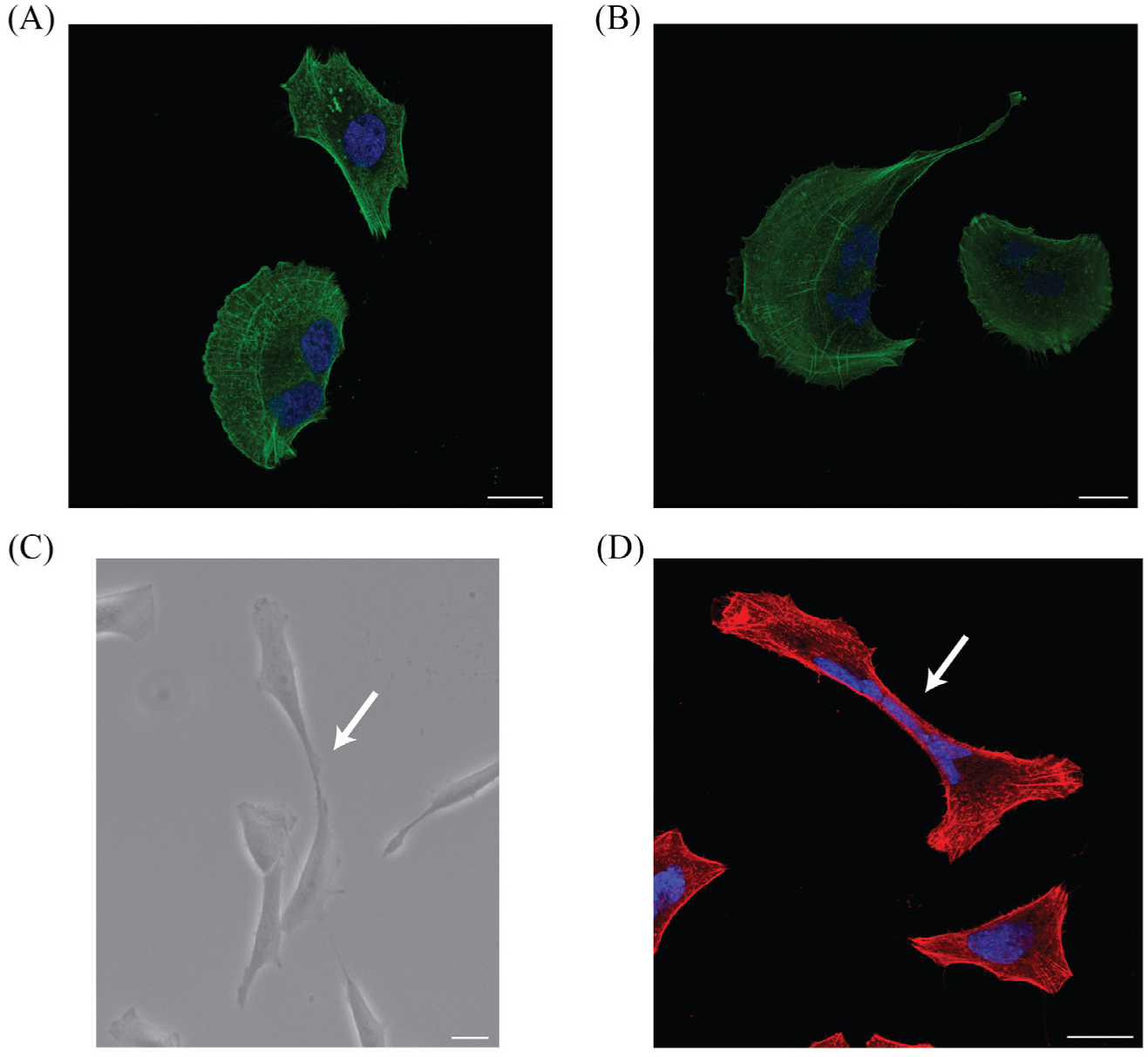
Disrupted cytokinesis in SEPT7 cKO Hs578T cells. (A,B) Immunocytochemistry of SEPT7 cKO cells stained for actin (green) and nuclei (blue) reveals the presence of giant multinucleated cells with an increased size (> 100 µm) and multiple nuclei, indicating a defect in cytokinesis. (C) Bright-field image of connected SEPT7 cKO cells (indicated by white arrow) with lamellipodia and leading edges in opposite directions. (D) Immunocytochemistry of SEPT7 cKO cells for actin (red) and nuclei (blue). Image shows a multi-nucleated giant cell that is tentatively in the process of dividing into two cells that crawl away from each other (indicated by white arrow). Scale bars are 25 µm.

**Figure S10.**
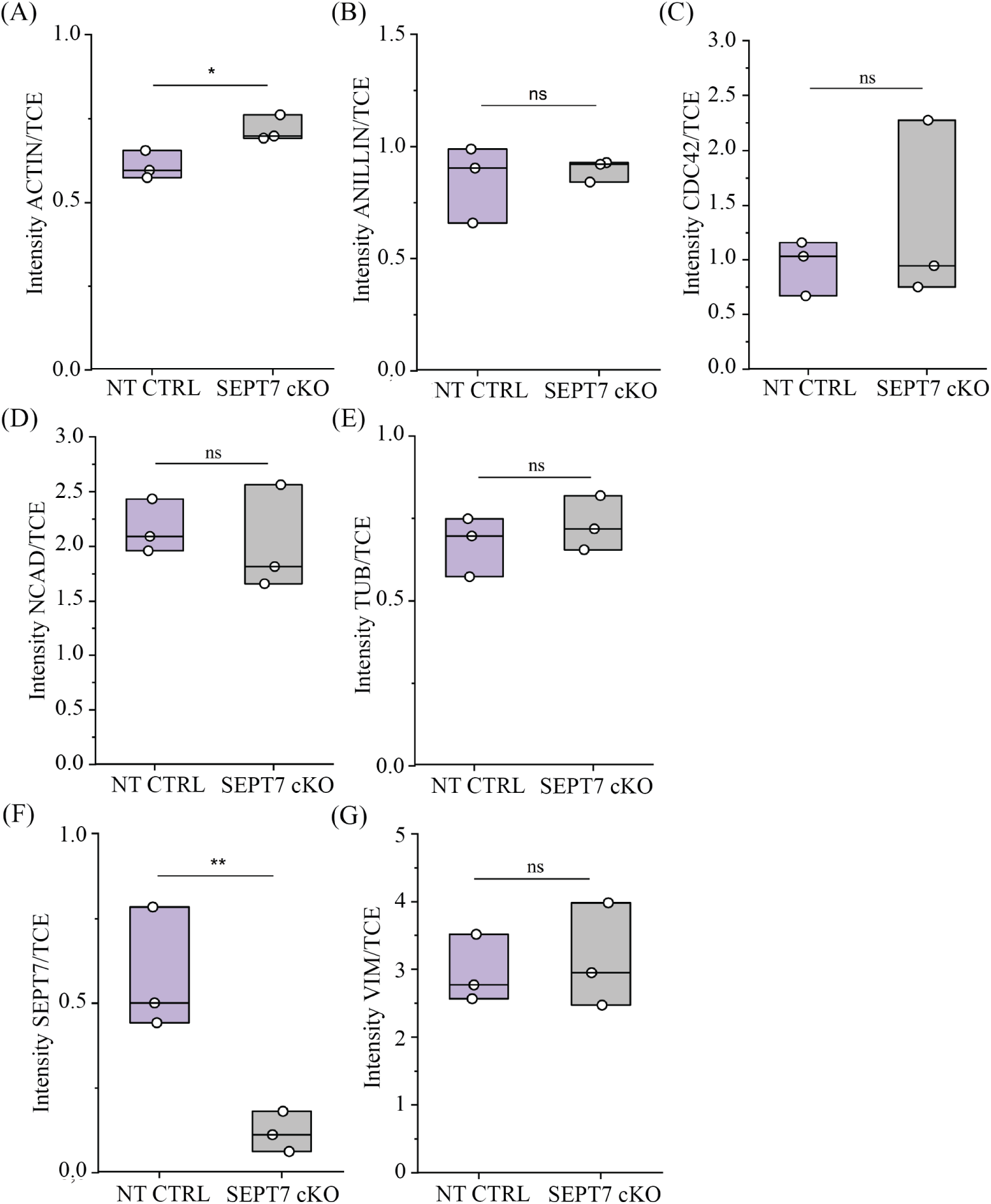
Western blot analysis of cytoskeletal for NT CTRL and SEPT7 cKO Hs578T cells. Protein levels are normalized by the TCE levels and compared for NT CTRL Hs578T (purple, left side) and SEPT7 cKO Hs578T (gray, right side). Data are shown for: (A) actin, (B) anillin, (C) CDC42, (D) N-cadherin, (E) α-tubulin, (F) SEPT7, and (G) vimentin. P-value results from t-tests are indicated by (ns) = p≥0.05, (*) = p<0.05, (**) = p<0.01. n=3 for each condition. We emphasize that the compared band intensities are always from the same blots, which are shown in their entirety in S13.

**Figure S11.**
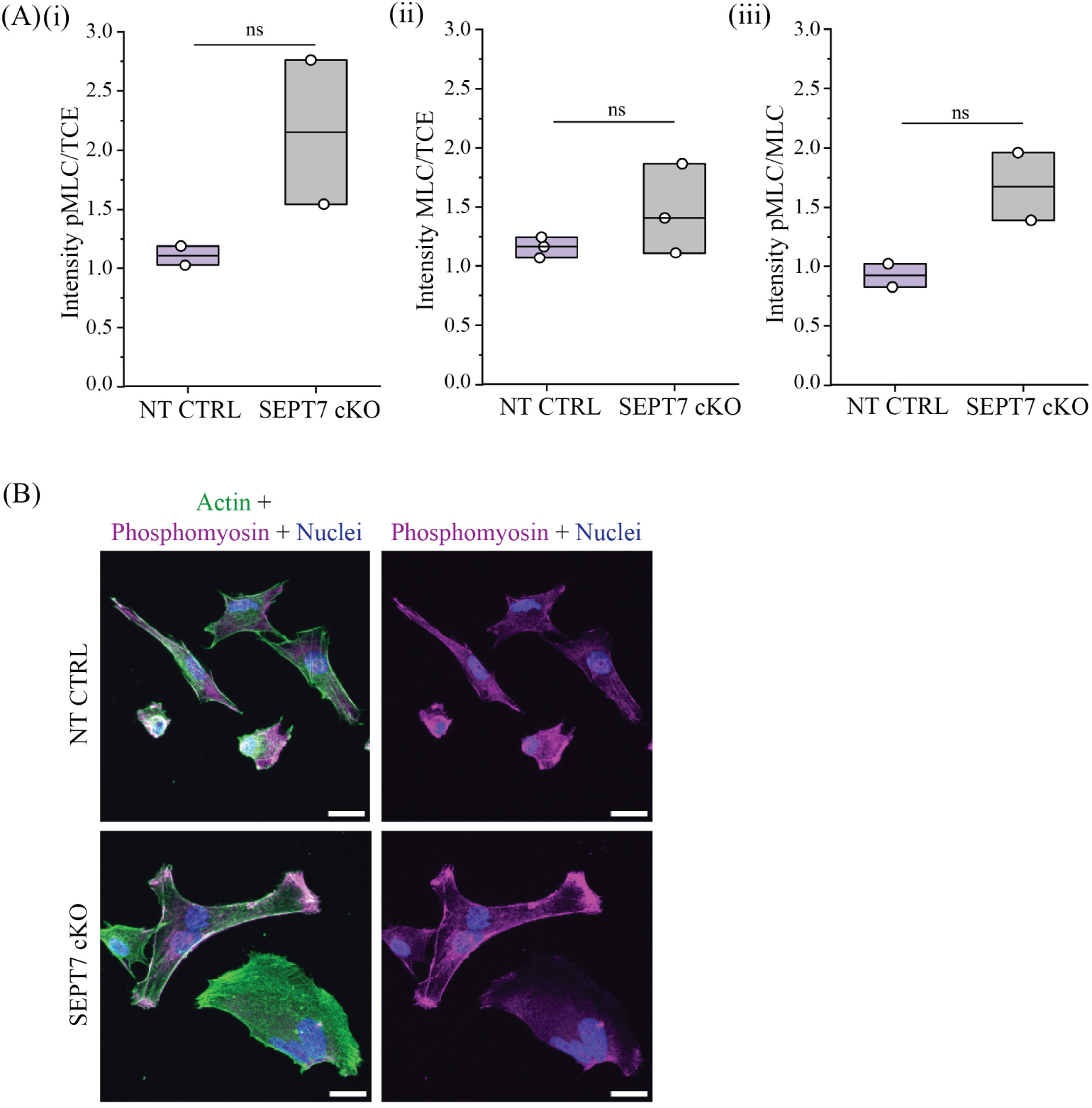
Western blot and immunocytochemistry analysis of nonmuscle myosin II activity in Hs578T cells. (A) Western Blot analysis of protein levels normalized by the TCE levels for NT CTRL Hs578T (purple, left side) and SEPT7 cKO Hs578T (gray, right side), for (i) phospho-myosin light chain (pMLC) and (ii) myosin light chain (MLC), and (iii) the ratio of pMLC/MLC. P-value results from t-tests are indicated by (ns) = p≥0.05. We emphasize that the compared band intensities are always from the same blots, which are shown in their entirety in S14. (B) Immunocytochemistry analysis of NT CTRL Hs578T (top row) and SEPT7 cKO (bottom row), for actin (green), phospho-myosin (purple) and nuclei (blue). Scale bars are 25 µM.

**Figure S12.**
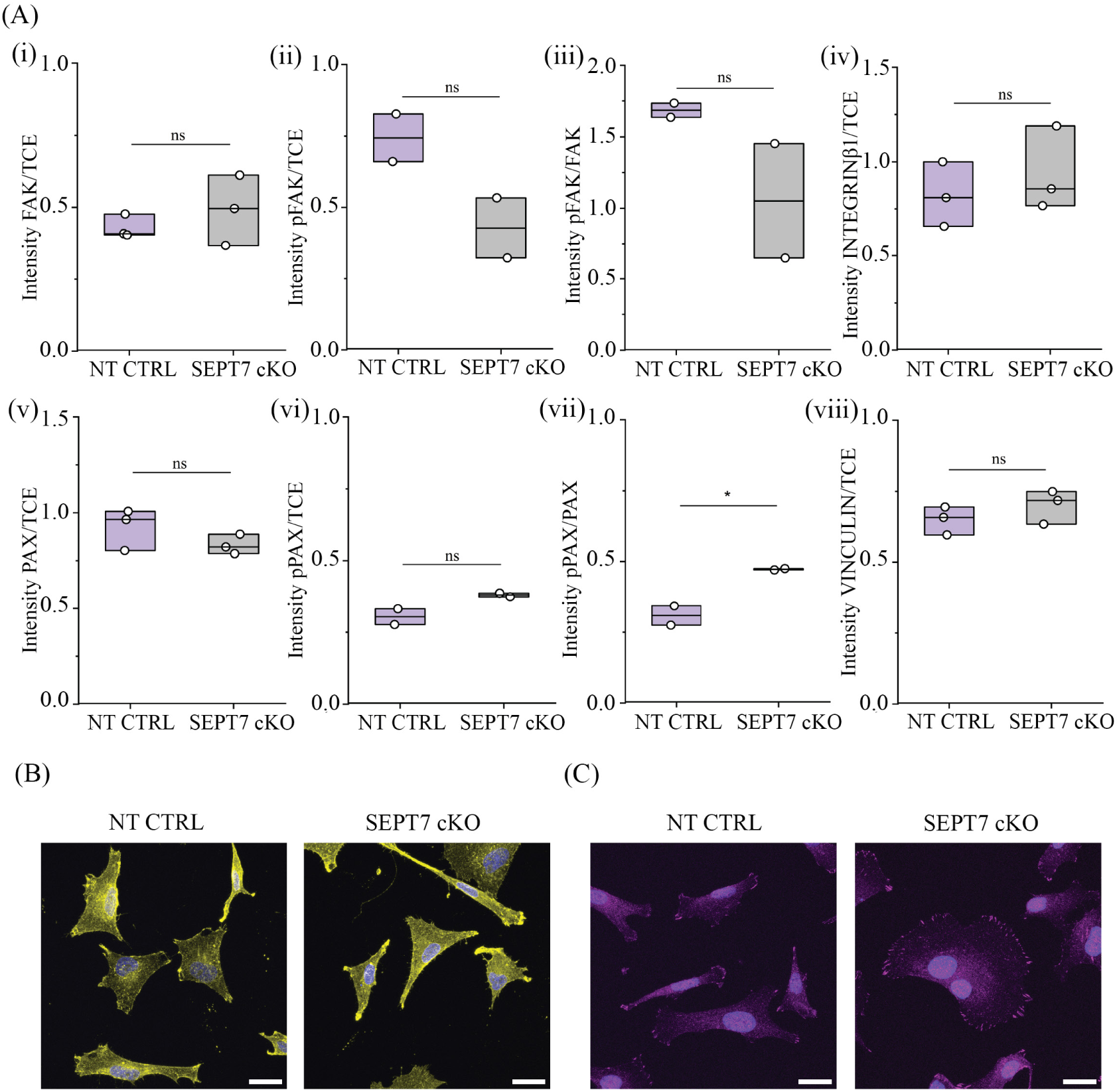
Western blot and immunocytochemistry analysis of focal adhesion proteins in Hs578T cells. (A) Western Blot analysis of protein levels normalized by the TCE levels for NT CTRL HS578T (purple, left side) and SEPT7 cKO Hs578T (gray, right side), for (i) focal adhesion kinase (FAK), (ii) phospho-focal adhesion kinase (pFAK), (iv) integrin-β1, (v) paxillin (PAX), (vi) phospho-paxillin (pPAX), and (viii) vinculin. Also shown are the ratios for (iii) pFAK/FAK and (vii) pPAX/PAX. P-value results from t-tests are indicated by (ns) = p≥0.05 and (*) = p<0.05. We hemphasize that the compared band intensities are always from the same blots, which are shown in their entirety in S15. (B, C) Immunocytochemistry analysis of NT CTRL (left side) and SEPT7 cKO (right side) cells for (B) integrin-β1 (yellow) and nuclei (blue), and (C) vinculin (purple) and nuclei (blue). Scale bars are 25 µM.

**Figure S13.**
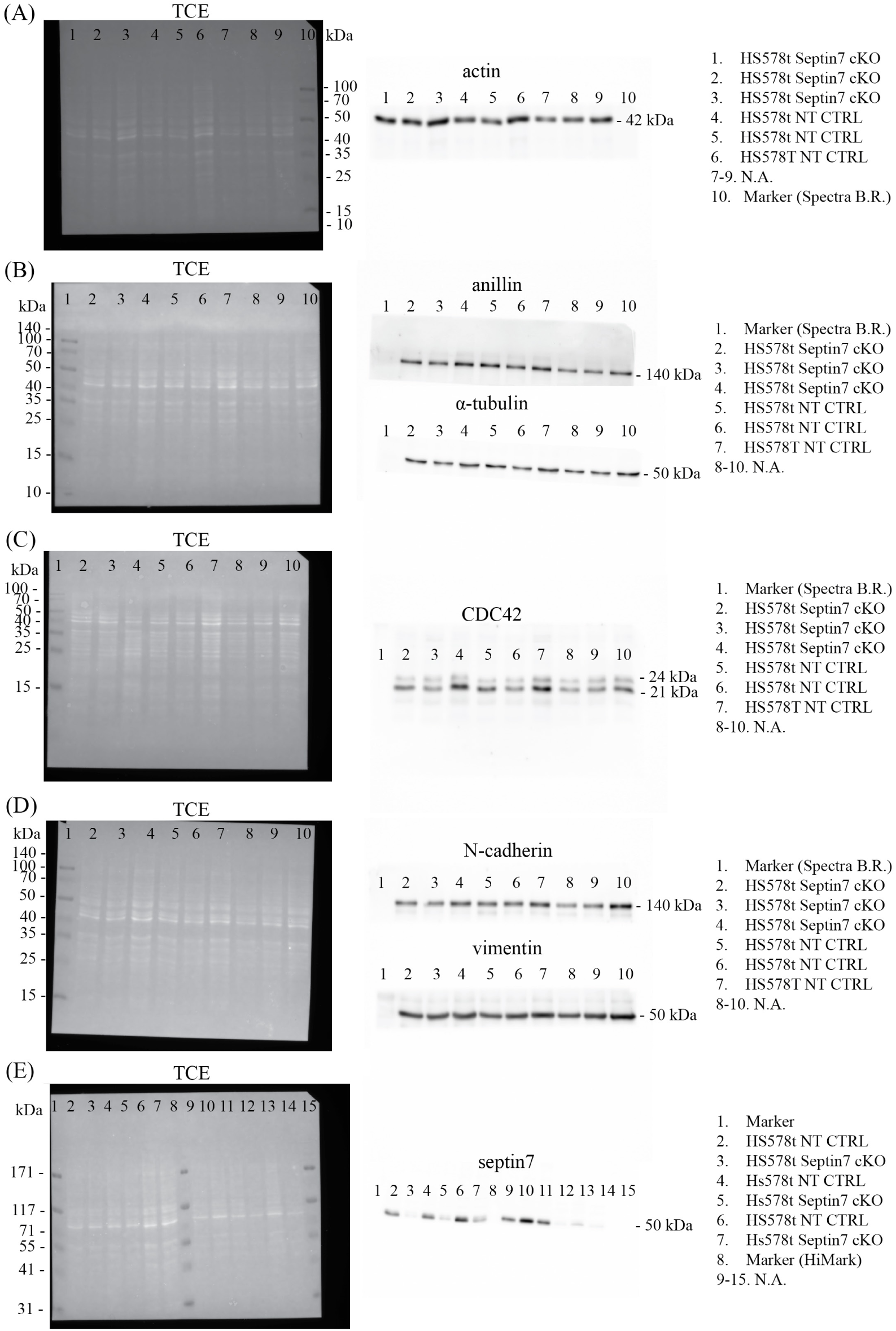
Full blots corresponding to Fig. S10. Shown are full TCE blots (left) and corresponding immunostained blots (center) with lane identification (right) for (A) actin, (B) annilin and α-tubulin, (C) CDC42, (D) N-cadherin and vimentin, and (E) septin 7.

**Figure S14.**
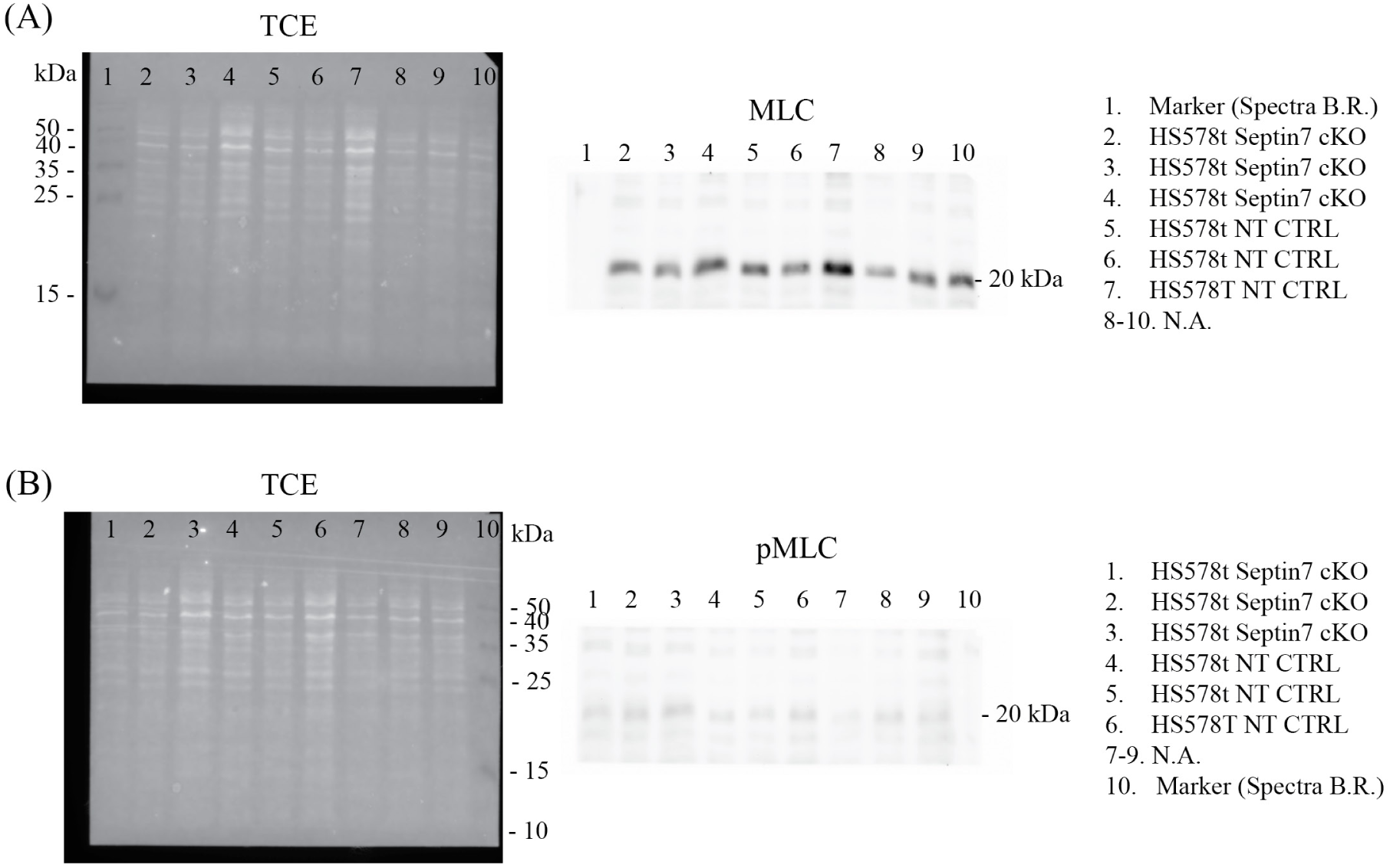
Full blots corresponding to Fig. S11. Shown are full TCE blots (left) and corresponding immunostained blots (center) with lane identification (right) for (A) MLC and (B) pMLC.

**Figure S15.**
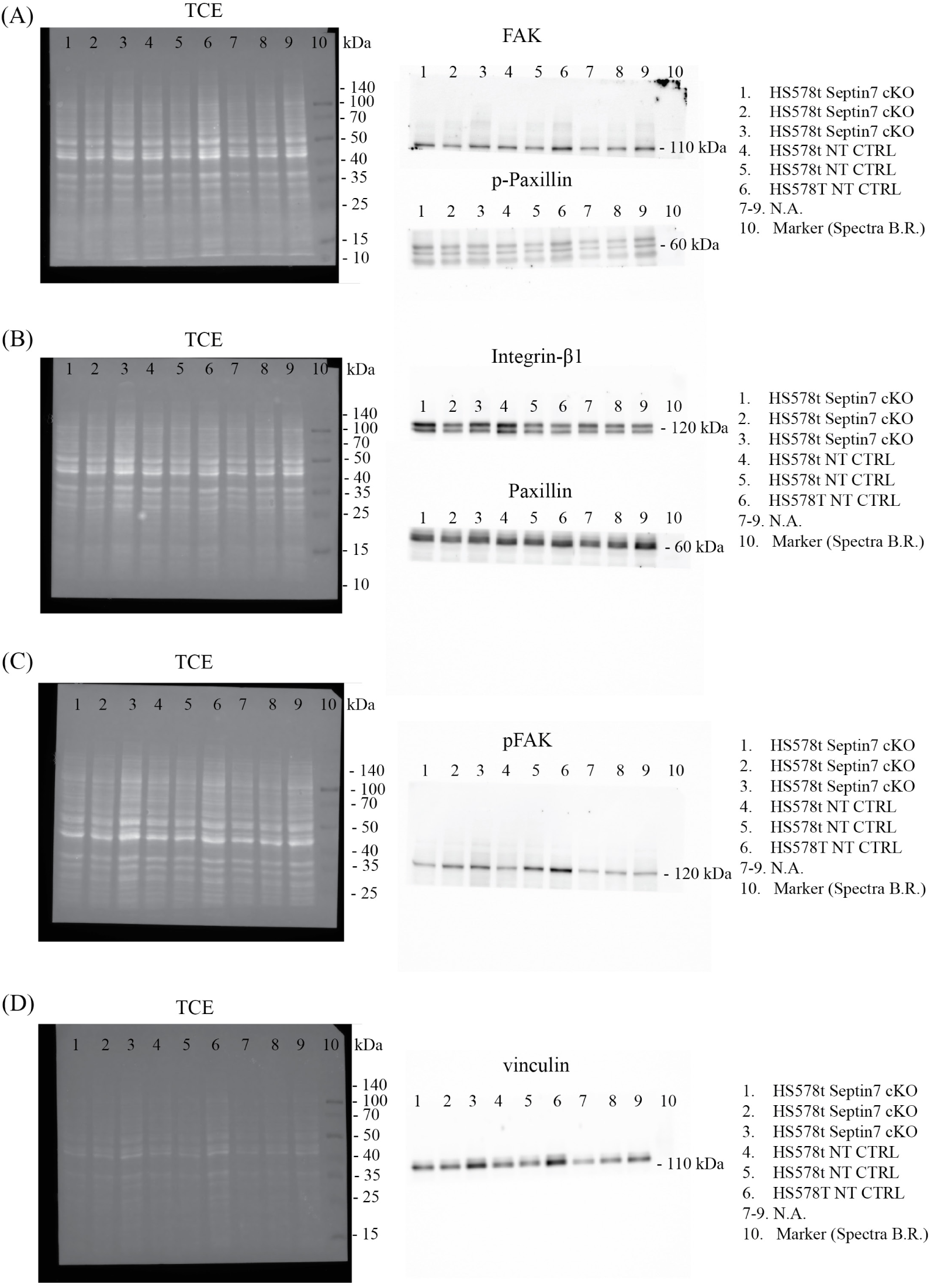
Full blots corresponding to Fig. S12. Shown are full TCE blots (left) and corresponding immunostained blots (center) with lane identification (right) for (A) FAK, p-paxillin, (B) integrin-β1 and paxillin, (C) pFAK, and (D) vinculin.

